# Chimeric and mutant CARD9 constructs enable analyses of conserved and diverged autoinhibition mechanisms in the CARD-CC protein family

**DOI:** 10.1101/2023.03.06.531260

**Authors:** Jens Staal, Yasmine Driege, Femke Van Gaever, Jill Steels, Rudi Beyaert

**Affiliations:** Unit of Molecular Signal Transduction in Inflammation, VIB-UGent Center for Inflammation Research, Ghent, Belgium; Department of Biomedical Molecular Biology, Ghent University, Ghent, Belgium; Department of Biochemistry and Microbiology, Ghent University, Ghent, Belgium

**Keywords:** protein engineering, signal transduction, inflammation, natural variation, UK biobank, reverse genetics

## Abstract

CARD9, -10, -11 and -14 all belong to the CARD-coiled coil (CC) protein family and originated from a single common ancestral protein early in vertebrate evolution. All four proteins form CARD-CC/BCL10/MALT1 (CBM) complexes leading to NF-κB activation after upstream phosphorylation by various protein kinase C (PKC) isoforms. CBM complex signaling is critical for innate and adaptive immunity, but aberrant activation can cause autoimmune or autoinflammatory diseases, or be oncogenic. CARD9 shows a superior auto-inhibition with very low spontaneous activity when overexpressed in HEK293T cells. In contrast, the poor auto-inhibition of other CARD-CC family proteins, especially CARD10 (CARMA3) and CARD14 (CARMA2), is hampering characterization of upstream activators or activating mutations in overexpression studies. We grafted different domains from CARD10, -11 and 14 on CARD9 to generate chimeric CARD9 backbones for functional characterization of activating mutants using NF-κB reporter gene activation in HEK293T cells as readout. CARD11 (CARMA1) activity was not further reduced by grafting on CARD9 backbones. The chimeric CARD9 approach was subsequently validated by using several known disease-associated mutations in CARD10 and CARD14, and additional screening allowed us to identify several novel activating natural variants in human CARD9 and CARD10. Using Genebass as a resource of exome-based disease association statistics, we found that activated alleles of CARD9 correlate with irritable bowel syndrome (IBS), constipation, osteoarthritis, fibromyalgia, insomnia, anxiety and depression, which can occur as comorbidities.

## Introduction

The CARD-CC/BCL10/MALT1-like paracaspase (CBM) complexes are critical for signaling leading to pro-inflammatory gene expression. Dysregulation of the CBM complexes has been associated to many immune-related diseases and cancer [1]. The CBM complexes are ancient and can be found as far back as Cnidaria [2–4]. In vertebrates, there are four distinct CBM complexes (CBM-9, CBM-1, CBM-2 and CBM-3) defined by the different CARD-CC protein family members: CARD9, CARD11 (CARMA1), CARD14 (CARMA2) and CARD10 (CARMA3), respectively [5–7]. This expansion of the monophyletic CARD-CC family coincides with an expansion of many functionally associated immune-related genes early in vertebrate evolution, most likely as a consequence of genome duplication [3,8,9]. CARD9 has retained the domain structure of the ancestral CARD-CC protein found in invertebrates, a sister group with the last common ancestor found in tunicates [10]. The other paralogs (CARD10, -11 and -14) have all obtained a C terminal fusion with a ZO-1/Dlg5-like MAGUK domain [2,10,11], forming the CARMA sub-family (Fig. 1, Fig. 2A). A common theme between the four jawed vertebrate CARD-CC proteins is that they all get activated by protein kinase C (PKC) [10], and will subsequently form filamentous polymeric filaments by aggregation via their CC domain [4,12–15]. These CARD-CC filaments will bind the signaling proteins BCL10 and MALT1 via the CARD domain, which subsequently will signal towards NF-κB activation and activation of the paracaspase MALT1 protease activity [16–21]. The recruitment of the BCL10/MALT1 complex to the CARD-CC protein is a critical signaling event and BCL10 fusion to CARD11 results in a constitutively active signaling complex [22]. The normal sequence of events is however most likely phosphorylation leading to CC domain aggregation and subsequent BCL10/MALT1 recruitment to the exposed CARD domain [15,23–26]. Despite their similarities, the four CARD-CC proteins are distinguished especially by their expression profile, where CARD9 and CARD11 are primarily expressed in innate and adaptive immune cells, respectively, and CARD10 and CARD14 primarily in different non-hematopoietic cells. The cell type specific expression differences also influence the phenotypes of mutants in the different CARD-CC family members [27]. Loss of function mutations in CARD11 cause severe combined immunodeficiency (SCID) due to a defective adaptive immunity [28]. Weaker hypomorphic mutations of CARD11, in contrast, cause hyper-IgE syndrome (HIES) with atopic dermatitis, a phenotype also seen when downstream MALT1 is lost [29–32]. Loss of function mutations in CARD9, on the other hand, cause a defective innate immune response to especially fungal infections [33,34], and the combined loss of CARD9 and MyD88 cause a severe disruption of innate immunity since two central adaptor proteins for different innate immune receptor signaling pathways are lost [35]. There are no strong reports on phenotypes from loss of function mutations in *CARD10* or *CARD14*, but it has been suggested that loss of CARD14 could sensitize to the development of atopic dermatitis [36]. Mice deficient for *Card10* show neurodevelopmental defects, a phenotype shared with *Bcl10* deficient mice [37,38]. Human germ line and somatic mutations in *CARD10* have been suggested to be associated to immunodeficiency, autoimmunity, cancer, cognitive defects and primary open-angle glaucoma (POAG), but it is not established whether the mutations are activating or inactivating [39–41]. High expression of CARD10 or CARD14 have both been associated to increased cancer cell survival [42,43]. Germ line activating mutations in CARD-CC proteins have up until now only been found in CARD11, which result in B cell lymphomas and *B cell expansion with NF-κB and T cell anergy* (BENTA) [44,45], as well as CARD14, which result in CAPE (*CARD14* – Associated Papulosquamous Eruption) – a disease spectrum that includes psoriasis and *Pityriasis rubra pilaris* (PRP) [46,47]. For both CARD14 and CARD10, the functional characterization of activating mutations is often hampered by the very high basal activity during transient overexpression studies in HEK293T cells, which leads to no or poor enhancement of the relative NF-κB activation from the activating mutations compared to wild-type [10,19]. We generated chimeric proteins in order to exploit the superior auto-inhibition properties of CARD9 to functionally characterize mutations from the other CARD-CC family members. Our chimeric CARD-CC proteins are artificially made by transplantation and replacement of functionally conserved parts to generate a novel but fully functional protein. The functionally conserved paralog sequences that we use have diverged during at least 500 million years of independent evolution, since the split between CARD9 and the CARMA proteins occurred before the last common ancestor of hagfish and jawed vertebrates [10,48,49].

**Figure 1.**
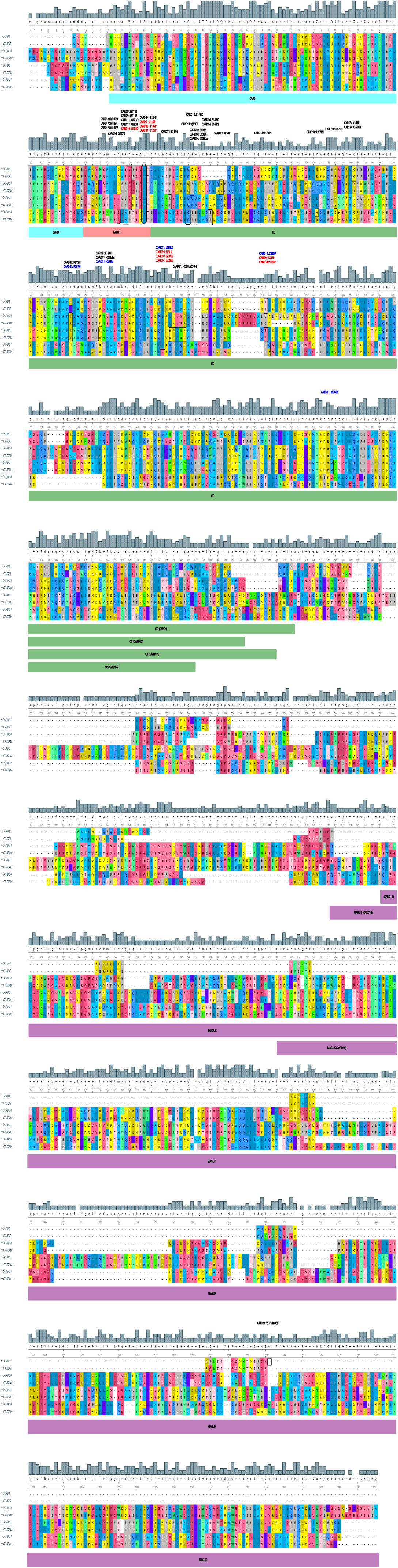
MUSCLE multiple sequence alignment of human and mouse CARD9 (human: NP_434700.2, mouse: NP_001032836.1), CARD10 (human: NP_055365.2, mouse: NP_570929.3), CARD11 (human: NP_001311210.1, mouse: NP_780571.2) and CARD14 (human: NP_001353314.1, mouse: XP_006532429.1) protein sequences. Domain start and end were determined by Uniprot domain annotations and manual graphical structure analysis of the Alphafold computed structural models. Activating natural variants (black), oncogenic somatic activating mutations (blue) and candidate artificial activating mutations (red) were designed based on homology indicated in the alignment.

**Figure 2.**
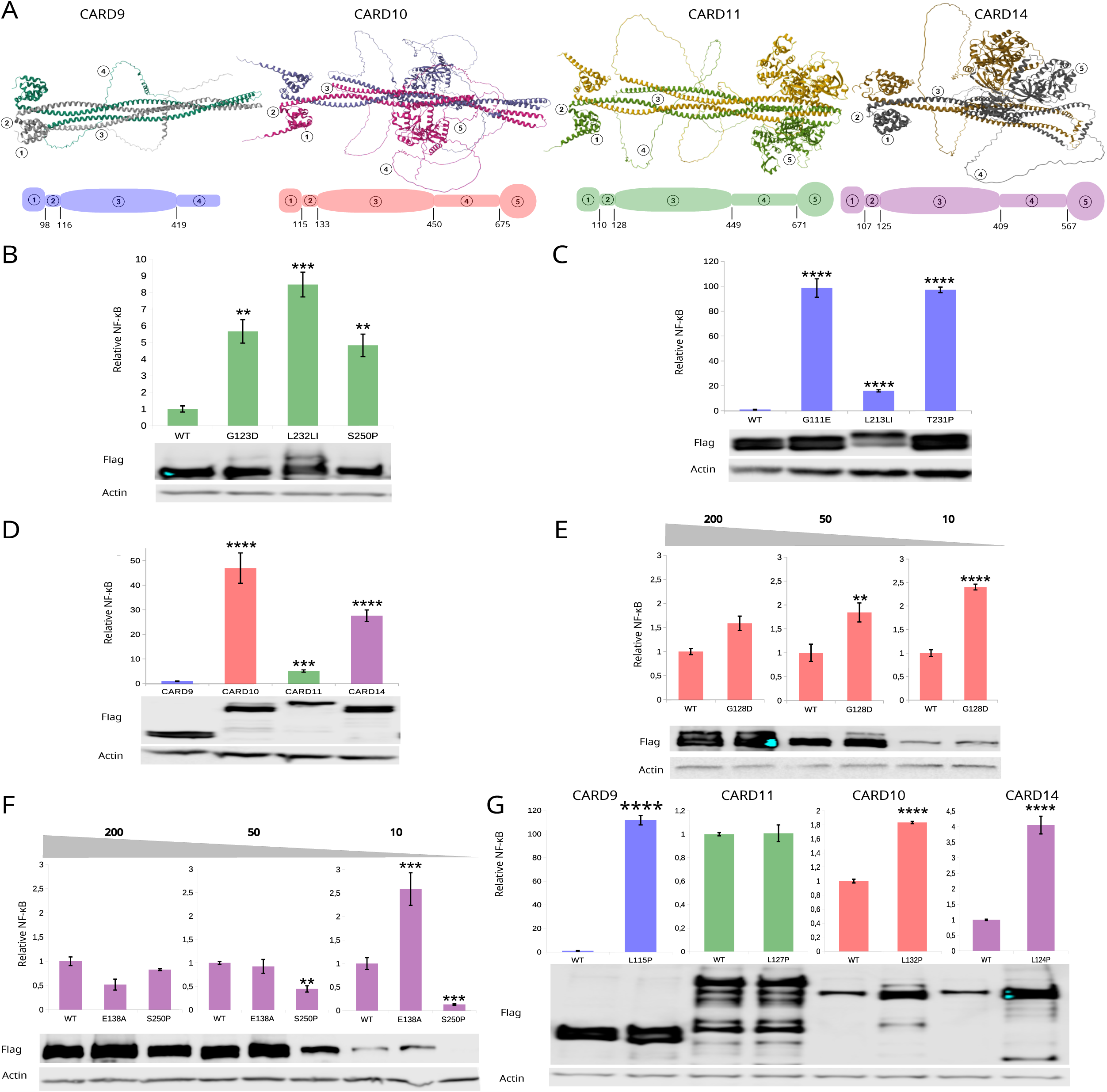
**A)** Comparison of the dimeric Alphafold-multimer computed structural models of the human CARD-CC family members CARD9, CARD10, CARD11 and CARD14 (same sequences as in the alignment in Fig. 1). Below: domain-composition cartoons with the colour codes used throughout the manuscript. The colour codes are the same as the ones used for the phylogenetic analysis in [10]. Domains: ① = CARD, ② = LATCH, ③ = CC, ④ = Linker / ID, ⑤ = MAGUK. **B)** NF-κB luciferase assay for known B-cell lymphoma or BENTA-associated activating mutations in CARD11 residues that are conserved in several CARD-CC family members. HEK293T cells were transfected with 200ng CARD-CC expression construct/μg transfection mix. Relative induction compared to WT CARD11. **C)** NF-κB luciferase assay for transferred activating CARD11 mutations in CARD9: G123D CARD9^G111E^ (CARD11^G123D^), CARD9^L213LI^ (CARD11^L232LI^) and CARD9^T231P^ (CARD11^S250P^). HEK293T cells were transfected with 200ng CARD-CC expression construct/μg transfection mix. Relative induction compared to WT CARD9. **D)** NF-κB luciferase assay for WT CARD9, CARD10, CARD11 and CARD14. HEK293T cells were transfected with 200ng CARD-CC expression construct/μg transfection mix. Relative induction compared to WT CARD9. **E)** NF-κB luciferase assay for WT CARD10 and an artificial activating mutation (G128D) corresponding to CARD11^G123D^. HEK293T cells were transfected in a gradient of 200, 50 and 10 ng CARD-CC expression construct/μg transfection mix. Relative induction compared to WT CARD10 per transfection concentration. **F)** NF-κB luciferase assay for WT CARD14, a psoriasis-associated activating mutation (E138A) and the potentially activating S250P mutation corresponding to CARD11^S250P^. HEK293T cells were transfected in a gradient of 200, 50 and 10 ng CARD-CC expression construct/μg transfection mix. Relative induction compared to WT CARD14 per transfection concentration. **G)** NF-κB luciferase assay for the highly conserved CARD14^L124P^ mutation transferred to all four CARD-CC paralogs. HEK293T cells were transfected with 200 ng CARD-CC expression construct/ μg transfection mix for CARD9 and CARD11, and 10 ng CARD-CC expression construct/ μg transfection mix for CARD10 and CARD14. Relative induction compared to the WT of each respective CARD-CC protein. The lower running band in CARD10 L132P in the Flag developed Western blot most likely corresponds to the 70kDa N-terminal MALT1-mediated cleavage fragment of CARD10 [87]. The lower running band in CARD14 L124P has not been reported before. The lower bands in the case of CARD11 might reflect degradation products that are visible due to the longer exposure times used to see CARD10 and CARD14 that were transfected at 20-fold lower DNA concentrations than CARD11. All luciferase experiments have been repeated at least 3 times with similar relative trends. The error bars represent the standard deviation from 3-4 technical replicates (different transfections). The bottom panels show the expression levels of the different constructs. Statistical analyses and probability levels expressed by stars (*) are described in Materials & Methods.

## Results

### Sequence alignment can identify activating mutations transferable between CARD-CC members

Despite approximately half a billion years of independent evolution, the four CARD-CC paralogs are very similar with conserved domains (Fig. 1; Fig. 2A). However, as illustrated by the Hamming dissimilarity matrix from the MUSCLE (MUltiple Sequence Comparison by Log-Expectation) sequence alignment [50,51] (Table 1A), the CARD-CC paralogs have diverged significantly compared to the relatively minor differences that can be seen between the mouse and human orthologs. As illustrated by both the multiple sequence alignment (Fig. 1) and the dissimilarity matrix (Table 1A) are the N-terminal sequences typically better conserved while the sequences downstream of the CC domain (Fig. 1; Table 1B) show poorer conservation. Previous structural comparisons between CARD9 and CARD11 have focused on the N terminal parts of the two proteins [15] and did not include the complete CC domain. In that structural comparison, they could determine that the CARD-LATCH bound to a small CC domain fragment formed a dimer. To investigate this further, we generated AlphaFold computed multimeric models [52] of CARD9, CARD10, CARD11 and CARD14 (Fig. 2A), which do seem to agree with the experimentally determined N-terminal structure [15]. The computed dimeric structural model of CARD9 overlaps with the experimentally determined dimeric N-terminal CARD9 structure with a root mean square deviation (RMSD) of 2.27 Å.

**Table 1.** The sequence difference between human and mouse orthologs is very small compared to the difference between the CARD-CC paralogs, which diverged prior to the evolution of sharks [2,10], as illustrated by the Hamming dissimilarity scores [50] after MUSCLE [51] alignment of the full-length proteins (Fig. 1) in UGene (A). We use human CARD9 and CARD14 and mouse CARD10 and CARD11 because these were available in the lab and previous work on different mutants has been done on these backbones. Coordinates of the different modules used for hybrid CARD-CC proteins from the selected proteins (B). Domain coordinates were taken from Uniprot and verified or complemented by (predicted) structure investigations using Alphafold and sequence alignments. MAGUK is defined as from the start of the PDZ domain.

|  | hCARD9 | mCARD9 | hCARD10 | mCARD10 | hCARD11 | mCARD11 | hCARD14 | mCARD14 |
| --- | --- | --- | --- | --- | --- | --- | --- | --- |
| hCARD9 | 0 | 72 | 340 | 327 | 308 | 307 | 375 | 378 |
| mCARD9 | 72 | 0 | 342 | 333 | 315 | 315 | 377 | 379 |
| hCARD10 | 340 | 342 | 0 | 85 | 627 | 631 | 666 | 687 |
| mCARD10 | 327 | 333 | 85 | 0 | 620 | 623 | 661 | 681 |
| hCARD11 | 308 | 315 | 627 | 620 | 0 | 91 | 672 | 703 |
| mCARD11 | 307 | 315 | 631 | 623 | 91 | 0 | 675 | 706 |
| hCARD14 | 375 | 377 | 666 | 661 | 672 | 675 | 0 | 234 |
| mCARD14 | 378 | 379 | 687 | 681 | 703 | 706 | 234 | 0 |

**B**
| # | Domain | hCARD9 |  | mCARD10 |  | mCARD11 |  | hCARD14 |  |
| --- | --- | --- | --- | --- | --- | --- | --- | --- | --- |
| ① | N-CARD | 1 [M...] | [...GKE] 98 | 1 [M...] | [...GQE] 115 | 1 [M...] | [...GKE] 110 | 1 [M...] | [...GLQ] 107 |
| ② | LATCH | 99 [PAR...] | [...GLT] 116 | 116 [PAQ...] | [...GLT] 133 | 111 [PTR...] | [...GLT] 128 | 108 [PDV...] | [...KLT] 125 |
| ③ | CC | 117 [QLL...] | [...QLE] 419 | 134 [QFL...] | [...TQG] 450 | 129 [HFL...] | [...QSL] 449 | 126 [ECL...] | [...QAE] 409 |
| ④ | Link/ID/Ct | 420 [TLV...] | [...*] 536 | 451 [GSC...] | 675 [...LLD] | 450 [PRH...] | [...PLV] 671 | 410 [PPG...] | [...ILS] 567 |
| ⑤ | MAGUK-C | N/A | N/A | 676 [SKA...] | [...*] 1021 | 672 [QHT...] | [...*] 1159 | 568 [QVT...] | [...*] 1004 |

CARD11 was the first CARD-CC family member where activating mutations were found, which cause spontaneous NF-κB activation and result in BENTA disease or B cell lymphoma [53]. Some of these activating mutations in CARD11 — for example CARD11^G123D^ [45,54], CARD11^L232LI^ [55,56], or CARD11^S250P^ [44] (Fig. 2B) — are mutated in residues that are conserved in several CARD-CC paralogs (Fig. 1). In structural comparisons of the N-terminal region of the CARD-CC proteins, it was demonstrated that several activating CARD11 mutations in the CARD and LATCH domains are translatable to CARD9 [15], for example the CARD9^G111S^ mutation, which corresponds to the CARD11^G123S^ mutation (rs387907352) that causes BENTA [57]. An interesting candidate germ line activating CARD9 mutation in the same residue is a natural CARD9 variant causing the very rare CARD9^G111E^ mutation (rs371695061). CARD9^G111E^ corresponds to the CARD11^G123D^ BENTA mutation (rs571517554) — which is more severe than CARD11^G123S^ [45]. By sequence alignment (Fig. 1), we can also see that CARD10^G128D^ would result in a similar artificial mutation in CARD10. We have already utilized the strategy of transferring the CARD11^L232LI^ oncogenic mutation located in the CC domain to CARD9^L213LI^ for functional characterization of an inactivating mutant [33]. The CARD9 T231 site is also highly interesting since it is critical for activation downstream of several different PKC isoforms [10,58]. The CARD9 T231 and CARD14 S250 residues align nicely with the oncogenic CARD11^S250P^ mutation, which means that CARD9^T231P^ and CARD14^S250P^ also are potential artificially activated mutations (Fig. 1). Interestingly, while the CARD9^L213LI^ gives a strong activation, this is clearly less than the exceptional activation that can be seen from the CARD9^T231P^ and CARD9^G111E^ mutations (Fig. 2C), demonstrating that these mutations are superior for functional characterization of activated CARD9. Consistent with previous observations [10,15,33], we see that inactive CARD9 shows reduced protein expression.

The CARD11^L232LI^ mutation has also previously been transferred to CARD10^L237LI^ and CARD14^L229LI^ (Fig. 1), but due to the high basal activation of the wild-type variants of these proteins (Fig. 2D), the relatively weak effect of these mutations was only apparent at very low expression levels [10]. This is however not unique for those mutations. For example, the activating artificial CARD10^G128D^ mutation (Fig 1) and the strong pustular psoriasis-associated CARD14^E138A^ mutation [59] do not show any visible enhancement of NF-κB activation at high expression levels, but do show a clear activating effect at lower expression levels (Fig 2E,F). Especially CARD14 appear sensitive to this overexpression effect. CARD14 exists in two different splice forms, and many previous analyses of CARD14 psoriasis-associated mutations have successfully used the less active, short splice form CARD14sh [60]. Surprisingly, in contrast to CARD11^S250P^ and CARD9^T231P^, the CARD14^S250P^ mutation was not activating, but rather detrimental for CARD14 activity. This may be due to the poor CARD14^S250P^ protein expression when low concentrations of the expression construct were transfected (Fig 2F). In contrast to CARD9, which has a strong autoinhibition even at high expression levels, the level of CARD14 expression will have an influence on NF-κB activation since WT CARD14 is spontaneously active when overexpressed in HEK293T cells (Fig. 2D). One possible reason for why CARD14^S250P^ fails to activate CARD14 is that the structural context of CARD9^T231^ and CARD11^S250^ is different from that of CARD14^S250^ (Fig. 2A). This serves as a nice illustration that primary sequence conservation alone is not a perfect predictor for translation of activating mutations between CARD-CC family members. CARD14 is in general less conserved in comparison to the other CARD-CC paralogs (Table 1A), and most psoriasis- and PRP-associated activating CARD14 mutation sites are not conserved in the other CARD-CC paralogs (Fig. 1). One exception is the PRP-associated mutation CARD14^L124P^ [61], which is highly conserved in all the paralogs (CARD9^L115^, CARD10^L132^, CARD11^L127^). The highly conserved CARD14 L124 residue sits at the end of the LATCH region, and the proline substitution probably disrupts a helix at the start of the CC domain (Fig. 1, Fig. 2A).Transfering this mutation to the other paralogs also results in the spontaneous activation of CARD9, -10 and 14, but surprisingly not CARD11 (Fig. 2G). This demonstrates that also known psoriasis or PRP-associated CARD14 variants in conserved residues can serve as a source of activating substitutions in the other CARD-CC family members.

### Identification of activating natural variants in CARD9

Since, to our knowledge, no germ-line activating mutations have previously been found or described in CARD9 [27], we investigated additional natural potentially activating substitutions from human sequence data in Uniprot [62]. At this moment, there is no disease phenotype associated with activated variants of CARD9. However, the top disease associations to CARD9^G111E^ (9-136371136-C-T) in Genebass [63] seem to be related to: constipation, IBS, fibromyalgia, arthritis and pain. Also behavioral issues seem to be associated to activated CARD9, such as insomnia, anxiety and depressive behaviors — often classified together in psychology as somatic symptom disorders (SSD). Interestingly, all of these disease conditions are well-known comorbidities [64–66], and a risk locus for IBS with constipation (IBS-C) in women has been mapped close to CARD9 on chromosome 9 [67]. Pain and misery (*dolor*) is one of the cardinal signs of inflammation [68], and it is thus possible that all these phenotypes originate directly or indirectly from activated CARD9. Also several other potentially activating mutations in addition to CARD9^G111E^ can be found in the human natural variation of CARD9, for example: L85F (rs1239721232; L85Y shown to be activating [15]), Y86C (rs1049019466; Y86F shown to be activating [15]), R101H (rs988002931; R101Q shown to be activating [15]) and V102I (rs1490862744). Testing these potentially activating natural variations in the LATCH region revealed that only CARD9^G111E^ caused strong spontaneous activation of CARD9 (Fig. 3A), with the other mutants being less active or as active as wild-type CARD9. The CARD9^G111E^ is as active as the very strong artificial CARD9^T231P^ mutation (Fig. 2C), indicating that it probably is close to the highest level of spontaneous activity that can be generated. One problem with CARD9^G111E^ is that it is an extremely rare mutation – with a single case in the Genebass database (out of 394841 individuals). The same mutation can not be found in gnomAD 2.1 [69], but G111R (9-139265589-C-T (GRCh37), rs769183409) has 3 individuals out of almost 250000 with an age range of 40-50 years old. This indicates that carriers of substitutions at this site are viable. There is no health data publicly available for these CARD9^G111R^ individuals, but gnomAD v2 typically only includes healthy individuals [70]. In the recently released gnomAD 4.0, which includes the UK biobank exome data, there are 6 individuals with the G111E variant and 9 with the G111R variant. Structural predictions at the G111 site indicate that any substitution at this site should be activating [15]. To test this, we also generated a CARD9 mutant (G111A) with a structural disruption as small as possible at this site. Interestingly, this substitution was almost as active as the G111E substitution (Fig. 3B), confirming the structural prediction that any substitution at G111 should be activating. Surprisingly is the G111R substitution much less active than the G111E and G111A substitutions. Different natural CARD9-activating mutations associated to the same sequelae would give more confidence to the predicted disease associations in Genebass, since it would exclude linkage to other disease-causing mutations in nearby genes. By listing missense mutations associated to one disease condition (fibromyalgia) associated to CARD9^G111E^ in Genebass, several other candidate natural mutations (K165E (rs781419489), S172N (rs201802044), E523Q (rs144943839) and *537Qext59* (rs1405807688)) can be identified which also have associations to psychological and gastrointestinal problems. By using alternative substitutions at the same site as an activating mutation, we can also identify additional potentially activating substitutions, such as K165del (9-136370972-CCTT-C) and E523K (rs144943839). Testing these mutations revealed that all of the fibromyalgia-associated variants and alternative variants in the same residues are activated, showing that fibromyalgia association is a very good predictor of activated CARD9 variants (Fig. 3B). Notably, the strongly activating variants in CARD9 were also predicted as pathogenic in AlphaMissense [71]. The S172N and E523Q variants are however showing weak (∼ 3-4 fold) activation. We consider these small differences less reliable than the big (qualitative) differences we typically see in activated CARD9 mutants. The L423V (9-136367639-G-C) mutation was not identified by association to fibromyalgia, but rather due to other comorbidities (constipation, pain) overlapping with the top disease associations for the K165E and *537Qext59* variants. The lack of activation in this variant highlights that fibromyalgia remains the best predictor of activated CARD9 variants for now. A full auto-activation survey of the CARD9 natural variations in humans and targeted investigations of CARD9 in patients with associated disease phenotypes for novel rare variants may be able to pick up even more disease-associated alleles in the future. Interestingly, a recently published association to the CARD9 R241Q mutation to osteoarthritis [72] may indicate that this mutation also is activating since osteoarthritis and fibromyalgia often are comorbid. It might also be possible to run iterative cycles of reverse genetics by *in vitro* testing of candidate mutations associated to the same diseases, and the new activated mutations could then form a new training set for another cycle of disease associations.

**Figure 3.**
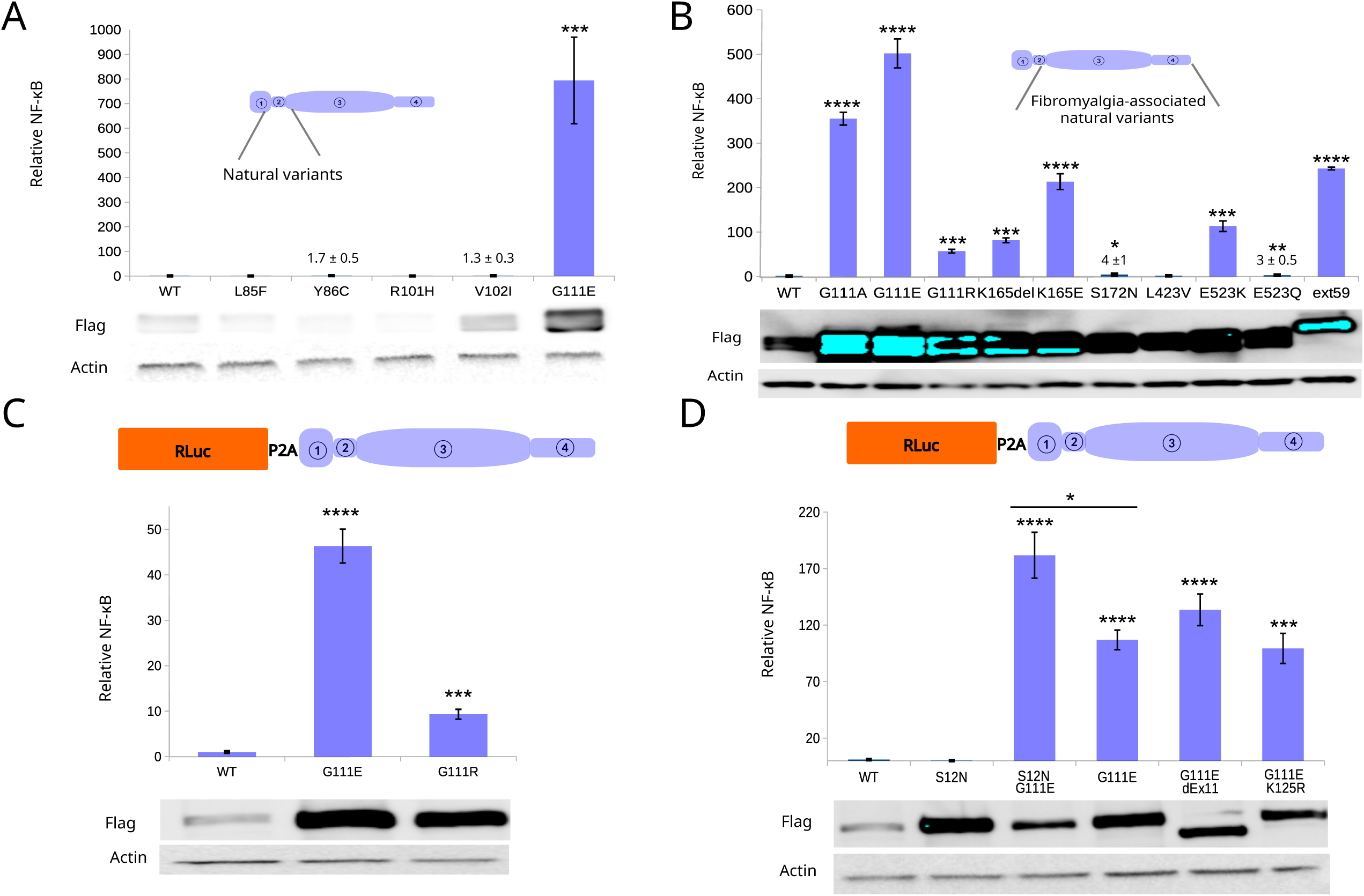
**A)** NF-κB luciferase assay for potentially activating natural variant mutations in CARD9 at the CARD-LATCH-CC interface transfected in HEK293T cells. Relative induction compared to WT CARD9. **B)** NF-κB luciferase assay of natural variants of CARD9 associated to fibromyalgia in Genebass, and alternative substitutions at the same residues, transfected in HEK293T cells. Relative induction compared to WT CARD9. Flag developed at high exposure to illustrate expression difference between WT CARD9 and the weak activating mutants. **C)** NF-κB luciferase assay of genetically activated CARD9^G111E^ and CARD9^G111R^ using RLuc-2A-CARD9 constructs driven by the CAG promoter, transfected in HEK293T cells at 100ng CARD-CC expression construct/μg transfection mix. **D)** Mechanistic studies genetically activated CARD9^G111E^ by NF-κB luciferase assay, using RLuc-2A-CARD9 constructs driven by the CAG promoter. HEK293T cells were transfected with 100ng CARD-CC expression construct/μg transfection mix. For (C) and (D): NF-κB firefly luciferase normalized against CARD9 mRNA expression via RLuc and NF-κB activation expressed as fold-induction compared to WT CARD9. All luciferase experiments have been repeated at least 3 times with similar relative trends. The error bars represent the standard deviation from 3-4 technical replicates (different transfections). The bottom panels show expression levels of the different constructs. Statistical analyses and probability levels expressed by stars (*) are described in Materials & Methods.

The screening for activated CARD9 variants also highlighted the problem with high expression differences between strongly activated variants and inactive variants or wild-type CARD9 (Fig. 3B). The CARD9 expressed in these experiments are on the pcDNA3 vector backbone and driven by the CMV promoter, which is known to respond to NF-κB and AP-1 [73,74]. The higher expression of activated CARD9 variants could thus be the result of a positive feedback loop. This positive feedback loop could however also be seen as a positive side-effect since it further enhances the contrast between inactive and activated variants.

### Characterization of activating natural variants in CARD9

The only potential “activating” disease-associated germ line CARD9 mutant known from literature would be CARD9^S12N^ [75], but this mutation is recessive and only enhances expression of some RelB-dependent cytokines after upstream receptor stimulation [75]. The enhanced RelB-dependent gene expression is supposedly due to reduced MALT1-mediated RelB cleavage [76], which means that the S12N mutation is not an activated CARD9 mutation. This mutation is thus more in line with other CARD11 mutations that show a complex phenotype with features of both gain- and loss-of-function [77]. Studies of the S12N mutant have revealed an exon 11 deletion (ΔEx11) rescue mutant [78], which abolishes the binding of TRIM62 to CARD9 [79]. The CARD9 ΔEx11 mutation leads to a frame shift mutation at the C terminal sequence, which has been shown to contribute to downstream signaling in activated CARD9 [79,80]. TRIM62, in turn, leads to K27-linked polyubiquitination at K125 on CARD9. Both of these events are critical for stimulation-induced CARD9 activation, since TRIM62 deficiency, CARD9 exon 11 deletion or the CARD9^K125R^ mutant lead to defective responses to upstream C type lectin receptor ligands [79]. To gain further mechanistic insight into whether genetically activated CARD9^G111E^ show similarities to stimulation-induced activation of CARD9, we re-cloned CARD9 in frame with *Renilla* luciferase (RLuc) and a P2A sequence for bicistronic expression of CARD9 and RLuc driven by a CAG promoter to avoid the potential feedback artifacts presented by the CMV promoter. Using RLuc, we can thus normalize NF-κB activity from the Firefly luciferase reporter to the number of CARD9 transcripts — which should lead to a specific measurement of the inherent difference in activity of CARD9. As a first verification, we tested CARD9^G111E^ and the less active CARD9^G111R^ (Fig. 3C). In line with the CMV-driven expression constructs, we could confirm the difference between the two different CARD9 G111 activating alleles and while the fold inductions were less dramatic due to the lack of a positive feedback loop, we still see very strong inductions from the strong activating CARD9^G111E^ mutant compared to WT CARD9 (Fig. 3C). Subsequently, we cloned CARD9^S12N^, and we also cloned the double mutants CARD9^S12N,G111E^, CARD9^G111E, ΔEx11^and CARD9^G111E, K125R^ for mechanistic investigations of genetically activated CARD9. As previously suggested [75], we could confirm that CARD9^S12N^ shows no significant elevation of basal activity compared to WT CARD9, but that the S12N mutation boosts already activated CARD9, as demonstrated by the CARD9^S12N,G111E^ double mutant (Fig. 3D). We could further establish that in contrast to stimulation-induced CARD9 activation, the activity of the genetically activated CARD9^G111E^ is independent of exon 11 or ubiquitination at the K125 site (Fig. 3D).

### The CARD9 C terminal sequence downstream of the CC domain is inhibitory in CARD9, -10, -11 and -14

Transfering activating mutations from CARD11 to CARD9 is generally far easier than transfering activating mutations from for example CARD14 to CARD9, since CARD11 has diverged far less from CARD9 and typically aligns better [2,10] (Fig. 1, Table 1A). In order to solve this issue, we investigated whether we could create hybrid proteins with the advantages of CARD9, such as the strong auto-inhibition and high fold-induction when activated, with sequence segments grafted from the other CARD-CC family members to enable direct transfer of known mutations into this chimeric CARD9 backbone. We have previously observed that deletion of the sequence downstream of the CC domain in CARD9, -10, -11 and -14 disrupts the auto-inhibition [2,33], indicating that the C-terminal part of CARD9 acts as the so-called “inhibitory domain” (ID) analogously to the linker sequence downstream of the CC domain in CARD11 [25]. Since this “ID” is an unstructured region and not a structurally defined or evolutionary conserved domain we will henceforth refer to it as the inhibitory region or linker. Also, in our screening for activated natural variants in CARD9, we identified two variants (E523K, *537Qext59*) that disrupt the autoinhibitory activity of this C-terminal sequence (Fig. 3B). With that in mind, we fused the CARD9 C-terminal sequence to the CARD-LATCH-CC part of CARD10, CARD11 and CARD14 (10N-9C, 11N-9C and 14N-9C). As comparison, we also generated proteins with a CARD9-like domain structure by deleting the C terminal MAGUK domain in the three CARMA proteins. While the CARD-CC segment of the protein family is well conserved, the linker / “inhibitory region” sequences are much less conserved (Fig. 1, Table 1A). Interestingly, despite the poor conservation, the CARD9 C-terminal inhibitory region sequence was either equally good or better than the native sequences downstream of the CC domains to provide some measure of auto-inhibition for all the CARD-CC variants (Fig. 4A). This is in contrast to previously generated hybrid proteins, where the CARD14 inhibitory region was unable to maintain CARD11 auto-inhibition [25].

**Figure 4.**
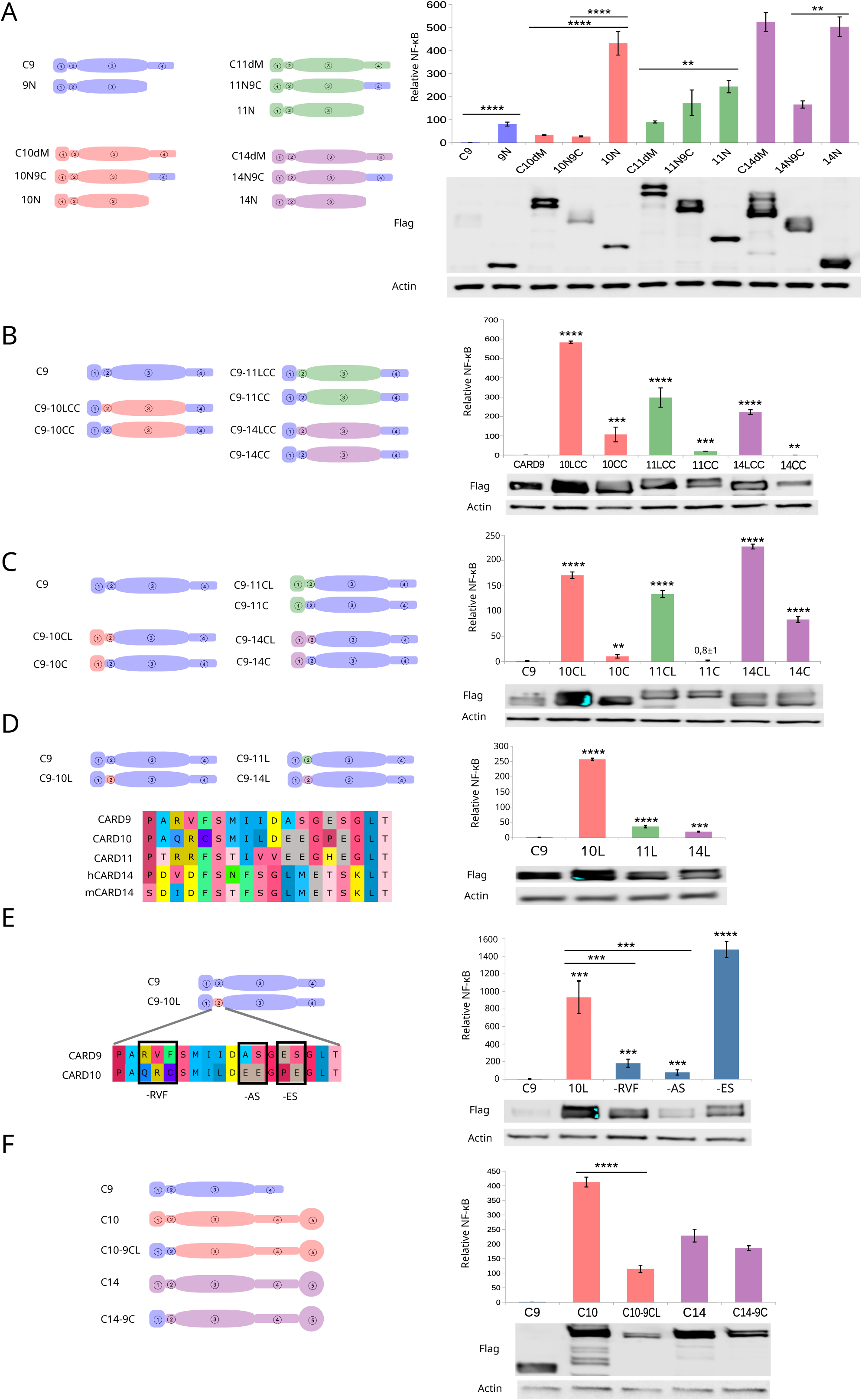
NF-κB dependent luciferase expression after overexpression of different chimeric CARD9 constructs in HEK293T cells. Relative induction is compared to WT CARD9 (C9) and protein expression levels analyzed with anti-flag via Western blotting. All luciferase experiments have been repeated at least 3 times with similar relative trends. The error bars represent the standard deviation from 3-4 technical replicates (different transfections). Statistical analyses and probability levels expressed by stars (*) are described in Materials & Methods. **A)** NF-κB luciferase assay for inhibitory activity of the CARD9 C-terminal sequence on CARD9, and the CARD-LATCH-CC fragments of CARD10, CARD11 and CARD14. Relative induction compared to WT CARD9. **B)** Comparisons of CARD9 and CARD9 with the LATCH-CC or CC domain replaced with the sequence from CARD10, CARD11 or CARD14. **C)** Comparisons of CARD9 and CARD9 with the CARD or CARD-LATCH domain replaced with the sequence from CARD10, CARD11 or CARD14. **D)** Comparisons of CARD9 and CARD9 with the LATCH domain replaced with the sequence from CARD10, CARD11 or CARD14. Multiple sequence alignment of CARD9, -10, -11 and -14 LATCH domains. Mouse and human LATCH sequence is identical for CARD9, -10 and -11, but not for CARD14. **E)** Effects of specific substitutions between CARD9 and CARD10 LATCH domain on auto-inhibition. **F)** CARD9 CARD transplanted to CARD14 and CARD9 CARD-LATCH transplanted to CARD10 compared to the wild-type CARMA proteins.

### The CARD9 CARD domain offers additional auto-inhibition

In a recent study focusing on the structure of the CARD, LATCH and early part of the CC domains, some residues were identified in the CARD9 CARD domain, compared to the CARD11 CARD domain, which contributed to an enhanced auto-inhibition of CARD9 [15]. We could see a similar effect from the C terminal deletion constructs, where the CARD9 N terminal construct showed less activity than the other corresponding constructs (Fig. 4A). Because of this, we replaced the CARD domain with the CARD9 CARD domain in our CARD-LATCH-CC clones with C terminal CARD9 (10N-9C, 11N-9C and 14N-9C), leading to a series of CARD9 LATCH-CC chimeras (CARD9-10LCC, CARD9-11LCC and CARD9-14LCC) (Fig 4B). The very strong spontaneous activation seen in the CARD14 clones (Fig 4A) was eliminated in the CARD replacements (Fig. 4B), indicating that a major reason for the poor auto-inhibition of the CARD14 chimeras lies in the CARD domain.

### CARD9 LATCH replacements influence auto-inhibition

Despite being a small sequence segment that is poorly conserved, especially in CARD14 [15], the LATCH (a.k.a α-linker) region between the CARD and CC domains is highly interesting since many activating mutations in CARD11 and CARD14 map to this region. Due to this, we initially also swapped the CARD-LATCH of 10N-9C, 11N-9C and 14N-9C to CARD9 (CARD9-10CC, CARD9-11CC and CARD9-14CC) leading to chimeras specific for the CC domains (Fig. 4B). While the CARD domain clearly influenced the spontaneous activity of CARD14-derived chimeras, spontaneous activity in CARD10-derived chimeras was specifically affected by the LATCH sequence. To ensure that this was a domain-intrinsic effect and not a specific interaction with the CC domain, we also did the opposite domain swapping and swapped either the CARD (CARD9-10C, CARD9-11C and CARD9-14C) or CARD-LATCH (CARD9-10CL, CARD9-11CL and CARD9-14CL) domain of CARD9 with the corresponding sequences from CARD10, CARD11 or CARD14 (Fig. 4C). This inverted effect verified that the small LATCH domain has a significant influence on the basal activity of the protein, which is corresponding to the location of many known activating mutations. It also verified that a major contributor to the poor auto-inhibition in CARD14-derived chimeras is the CARD domain. That the effect was visible independently of which CC domain was used indicates that the effect is not entirely due to specific inter-domain interactions as is proposed by the standard model of CARD-CC activation [56]. Inter-domain interactions do however have some effect since we see synergistic effects from CARD and LATCH domains in spontaneous background activity (Fig. 4B,C). To study the LATCH domains in isolation, we also replaced only the LATCH (CARD9-10L, CARD9-11L and CARD9-14L) domains (Fig. 4D), which strongly highlighted and verified the role of the CARD10 LATCH domain in the poor auto-inhibition. Based on sequence comparisons, it is very surprising that CARD10 and CARD11 LATCH domains cause less auto-inhibition than the CARD14 LATCH, which is a poorly conserved sequence (Fig. 4D). The CARD10 LATCH only has 7 significant substitutions in 3 segments, compared to the CARD9 LATCH (Fig. 4D,E). We found that two of the three substitution segments between the CARD9 and CARD10 LATCH domains contribute additively to the auto-inhibition by reverting the CARD10 LATCH sequence to the corresponding CARD9 sequence in each segment (CARD9-10L-RVF (=QRC/RVF substitution), CARD9-10L-AS (=EE/AS substitution), CARD9-10L-ES (=PE/ES substitution)) (Fig. 4E), which means that most of the CARD9 LATCH is needed for lower background signal in CARD10-derived chimeric CARD9 constructs. In order to validate that the mapped domains for auto-inhibition also are relevant in the original backbone, we generated a CARD10 clone with the CARD-LATCH transplanted from CARD9 (CARD10-9CL) and a CARD14 clone with the CARD transplanted from CARD9 (CARD14-9C). While the LATCH clearly is contributing to the auto-inhibition in CARD14 (Fig. 4B,C), it also is a site where many psoriasis-associated mutations are located (Fig. 1). Compared to wild-type CARD10, CARD10-9CL was clearly less auto-activated (Fig. 4F), which means that this could be a good backbone for mutant testing since most disease-associated mutations in CARD10 are located in the CC domain or further downstream [40]. CARD14 with the CARD domain replaced with CARD9 CARD did however not show any notable reduction of auto-activation compared to wild-type CARD14 (Fig. 4F), indicating that more domains are important in CARD14 regulation.

### Using the chimeric CARD9 constructs to analyze activating mutations in other CARD-CC proteins

Since CARD9 shows a superior auto-inhibition compared to the other CARD-CC proteins (Fig 2D) is the utilization of chimeric CARD9 constructs (Fig. 4) to study activating mutations in non-conserved residues an attractive idea. A low background activity should result in a high fold-induction and high signal-to-noise method for evaluation of activating mutations. As two case studies, we use the B cell lymphoma CARD11 CC mutation L232LI [55] and the pustular psoriasis CARD14 CC mutation E138A [59]. For the CARD11 L232LI mutation, we could see an interesting inter-domain interaction effect: while the corresponding CARD9^L213LI^ mutation was activating, the CARD9 with only CARD11 CC was not activated by the L232LI mutation (Fig. 5A). Activation by the CARD11^L232LI^ mutation required CARD11 LATCH. Taken together, this indicates that activation by the L/LI substitution requires a matching LATCH-CC interaction. In contrast, the activation by the CARD14^E138A^ mutation was domain-intrinsic and the strongest relative fold-induction was seen in CARD9 with only the CARD14 CC domain transplanted (Fig. 5B). While the CARD9-14LCC (CARD14 LATCH-CC) showed less than 2-fold activation from the E138A mutation, the same mutation in the CARD9-14CC backbone caused several 100-fold activation, demonstrating the importance of low background activity to reliably detect the effect of activating mutations by transient overexpression.

**Figure 5.**
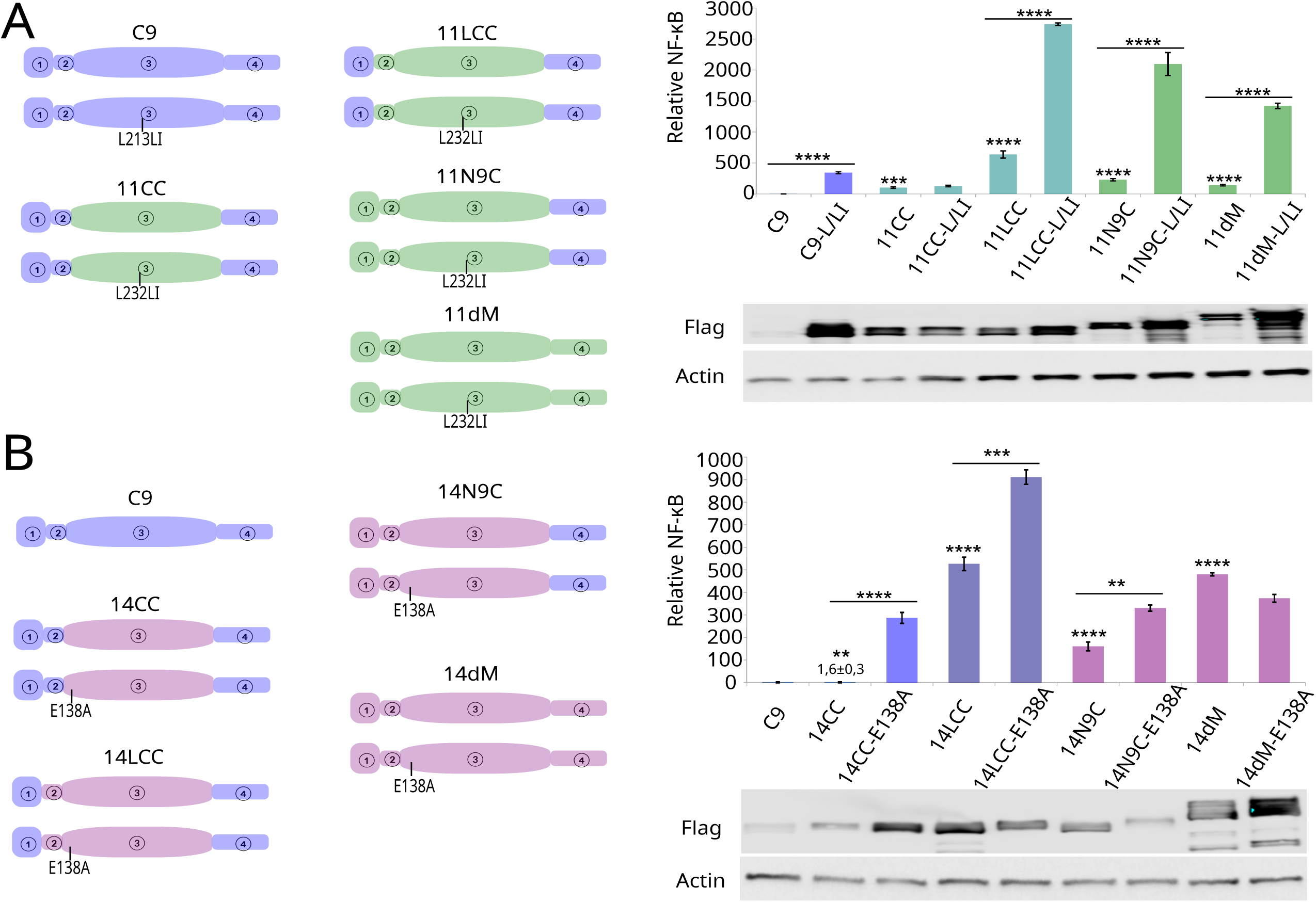
NF-κB dependent luciferase expression after overexpression of different chimeric CARD9 WT or activated mutant constructs. Relative induction compared to WT CARD9 (C9). Bottom panels show protein expression levels analyzed with anti-flag via Western blotting. All luciferase experiments have been repeated at least 3 times with similar relative trends. The error bars represent the standard deviation from 3-4 technical replicates (different transfections). Statistical analyses and probability levels expressed by stars (*) are described in Materials & Methods. **A)** Comparison of the effect of the L232LI mutation in CARD11 in different chimeric CARD9 backbones, compared to WT and the L213LI mutation in CARD9 **B)** Effect of the CARD14 psoriasis-associated mutation E138A in different chimeric CARD9 backbones, compared to corresponding wild-type constructs.

Since the CARD14 LATCH domain and the CC domains from CARD10 and CARD14 caused relatively mild auto-activation after transplantation to CARD9 (Fig. 4B,D), we investigated published mutations in these domains as a proof-of-concept of the chimeric CARD9 strategy. For CARD14 LATCH we could not see any activation from LATCH domain psoriasis-associated mutations, except for the highly conserved PRP mutation L124P [61] (Fig. 6A). This surprising result indicates that the activation from some CARD14 LATCH mutations, for example at positions G117 and M119, require specific interactions with other CARD14 domains. Such interactions are probably at the end of CARD and beginning of CC [15], but the computed structural models do not give any indications of such direct interactions (Fig. 2A). This does challenge the model that activating mutations are disrupting inter-domain interactions, however, since the CARD9-14L chimera is auto-inhibited despite the poor conservation of the CARD14 LATCH (Fig. 1, Fig. 4D). We had already seen that CARD9-14CC was a good candidate chimeric CARD9 construct (Fig. 5B), and this allowed us to investigate several psoriasis and PRP-associated mutations in the CARD14 CC domain to confirm that this strategy is generally applicable. All the mutations were strongly activating in the CARD9-14CC construct (Fig. 6B). The E138 allelic series showed that the PRP mutation E138K was a stronger activator than the two psoriasis-associated mutations E138A and E138del, in line with the suggested correlation between mutation strength and disease phenotype [46].

**Figure 6.**
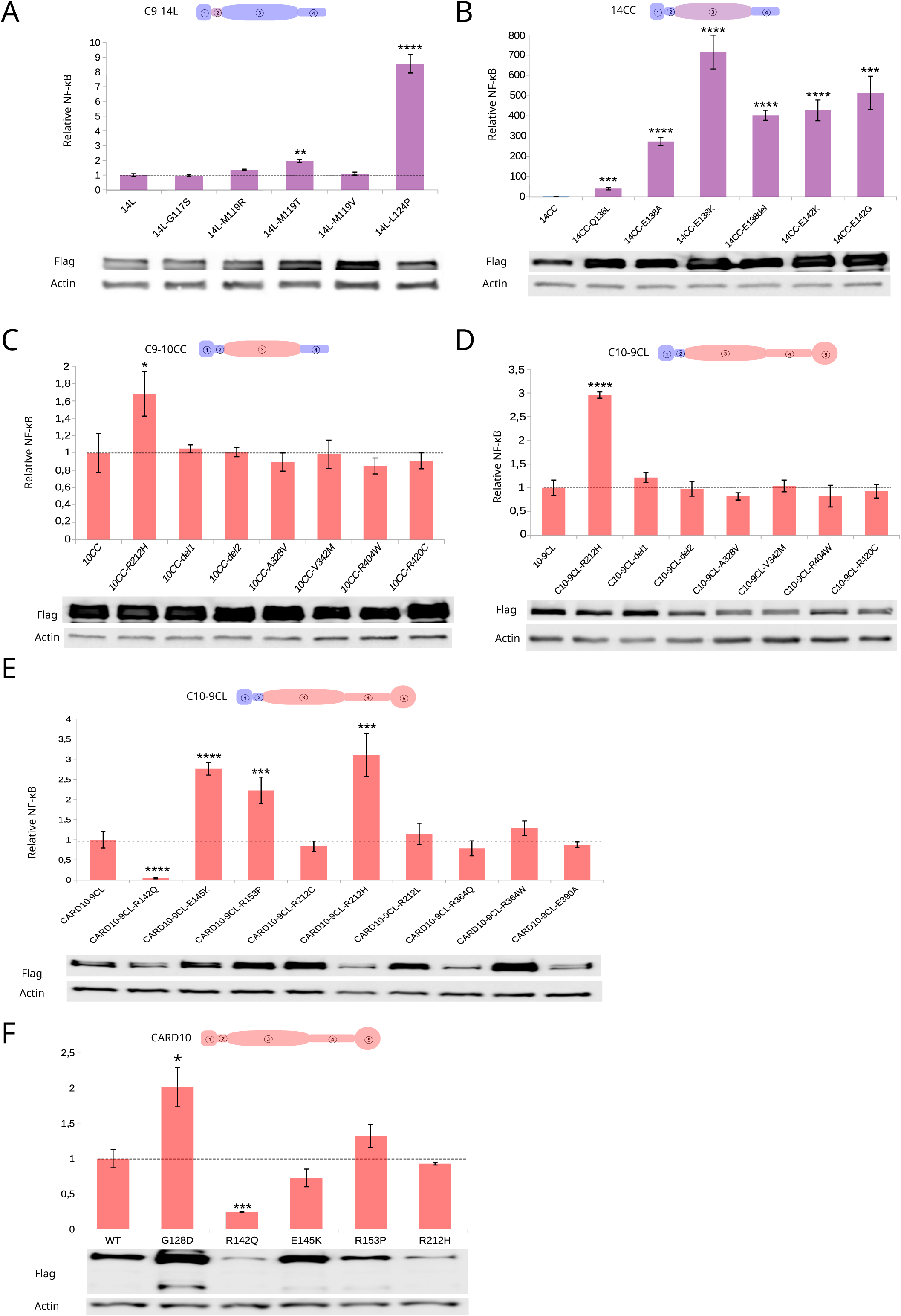
Proof-of-concept analysis of published mutations in the LATCH-CC region in CARD10 and CARD14, using the minimal chimeric CARD9 backbones with either the CARD9 LATCH or CC domain replaced with the corresponding sequence from CARD10 or CARD14. Only some mutations have previously been verified *in vitro* to be activating. Mutations are evaluated as relative NF-κB dependent luciferase expression after overexpression of different chimeric CARD9 constructs, compared to the wild-type chimeric CARD9. All luciferase experiments have been repeated at least 3 times with similar relative trends. The error bars represent the standard deviation from 3-4 technical replicates (different transfections). Statistical analyses and probability levels expressed by stars (*) are described in Materials & Methods. **A)** Evaluation of CARD14 LATCH mutations associated to psoriasis in the CARD9-14L backbone. **B)** Evaluation of CARD14 CC mutations associated to psoriasis in the CARD9-14CC backbone. **C)** Evaluation of CARD10 CC mutations associated to POAG, cancer or autoimmune immunodeficiency in the CARD9-10CC backbone. **D)** Evaluation of CARD10 CC mutations associated to POAG, cancer or autoimmune immunodeficiency in the CARD10-9CL backbone. **E)** Evaluation of CARD10 natural variants with various Genebass disease associations overlapping with the associations found for R212H in the CARD10-9CL backbone. Dashed line represents the normalized average NF-κB induction level of WT CARD10-9CL. **F)** Evaluation of activating CARD10 mutations in a pure CARD10 backbone. Dashed line represents the normalized average NF-κB induction level of WT CARD10. The lower running Western blot band in the case of G128D, E145K and R153P most likely corresponds to the 70kDa N-terminal MALT1-mediated cleavage fragment of CARD10 [90].

At this moment, no association has been made between a disease condition and activating mutations in CARD10 [27]. However, several germline mutations in the CARD10 CC domain have been found associated to diseases, such as primary open angle glaucoma (POAG), or as a single case of autoimmunity / immunodeficiency (R420C) [39,40]. There are also somatic CARD10 mutations described in cancers [41], and since CARD10 overexpression is associated to some cancers [81], those mutations are also good candidates for activating CARD10 mutations. Surprisingly, all mutations except one (R212H; ClinVar: VCV000224912.1; dbSNP: rs1057519378) were as active as the wild-type CARD9-10CC construct (Fig. 6C). Interestingly, CARD10 R212 aligns with R207 in CARD11 — and the CARD11 R207H mutation has been picked up in genetic B cell lymphoma screenings [82,83] — which supports that this mutation is likely to be activating. In comparison to the extremely rare activated CARD9^G111E^ allele is CARD10^R212H^ far more common, with 25 cases out of 394841 being reported in the Genebass database. The lack of activation from the other CARD10 mutations tested could either be a limitation of the chimeric CARD9 construct due to inter-domain dependencies similar to the CARD11 L/LI mutation (Fig. 5A), or biologically true since there is no prior evidence that these disease-associated mutations are activating. To investigate the first possibility, we re-cloned the mutations into the CARD10-9CL chimeric CARD10 construct, which shows much lower auto-activation compared to wild-type CARD10 (Fig. 4F). In line with the CARD9-10CC constructs, R212H was the only CARD10 mutation that activates also in the CARD10-9CL backbone (Fig. 6D). In this initial screening we have only evaluated CARD10 variants that have been associated to disease in literature. Since published disease associations seem to be a poor predictor of activating CARD10 variants, a broader screening was done. The R212 position has multiple natural variations (R212C (rs760871700), R212L (rs1057519378)), which potentially could represent additional activating variant alleles. Sorting for candidate activating mutations by disease in Genebass did not pick up candidates with an as uniform pattern of comorbidities in CARD10 as with activated CARD9, but some candidates (R142Q (rs199764326 ; cardiovascular), E145K (rs769979914), R153P (rs1187928736; kidney), and R364Q (rs199557110 ; allergic rhinitis)) were considered good starting points for seeding a more accurate prediction model for activating mutations based on disease associations. We included the E145K variant based on its similarity to the activating psoriasis-associated substitutions E138K and E142K in CARD14 (Fig. 6B). The screening verified two new activated variants (R153P, E145K), and surprisingly one inactivated variant (R142Q) (Fig. 6E). Based on disease associations in common between the verified activated variants available in Genebass (R153P, R212H), “acute nephritic syndrome” is the most promising disease association as a predictor of new candidate activated CARD10 variants. Using this association, we tested E390A (22-37507851-T-G) and R364W (rs184229951), which were not activating (Fig. 6E). This means that disease associations so far have failed to reliably predict activating CARD10 variants. In contrast to the psoriasis-associated CARD14 mutations in the CARD9-14L and CARD9-14CC backbone, the activating CARD10 mutations E145K, R153P and R212H only result in a very moderate relative activation. In order to evaluate their level of activation compared to the artificially activated CARD10^G128D^ as positive control, we also re-cloned these mutations in a pure CARD10 backbone. We also re-cloned the potentially inactivating mutation R142Q for validation on a pure CARD10 backbone. However, in a pure CARD10 backbone, none of the identified activating natural variants could be verified and only the strong artificially activated CARD10^G128D^ was significantly different from WT (Fig. 6F). The identified natural activating variants are thus most likely weak hypermorphic mutations, which get masked by the spontaneous activity of overexpressed CARD10. They may still show enhanced basal activity at endogenous expression levels or in a different cellular background, but this remains to be investigated. The surprising finding of an inactivating (R142Q) CARD10 mutation is also very interesting, since no human disease has been associated with CARD10 loss-of-function. All other CARD-CC members have reported diseases from loss-of-function in humans: CARD11 loss causes severe combined immunodeficiency (SCID) or atopic dermatitis in hypomorphic mutations, CARD9 loss causes impaired innate immunity against fungi and CARD14 loss has been suggested to cause atopic dermatitis. Screening for additional inactivating CARD10 natural variants could thus also be informative and lead to the identification of diseases associated to CARD10 loss-of-function in humans.

## Discussion

Regulation of CBM complex signaling is critical since dysregulation can cause inflammatory diseases [59] and cancer [55,84,85]. Transient transfection in HEK293T cells of signaling proteins is often used as a fast and simple approach to elucidate functional effects of different variants. Functional characterization of mutations in some of the CARD-CC proteins upon transfection in HEK293T cells can however be problematic due to their high base line activity, except for CARD9 that is strongly auto-inhibited [10,19,59]. A chimera is a highly useful concept and tool for functional characterization of highly divergent but functionally conserved parts [86]. Starting from the observation that overexpression of CARD9 is completely functionally silent, we have demonstrated that hybrid chimeric proteins that use CARD9 as a backbone enables studies of mutants from the other CARD-CC family members by swapping the corresponding region between both the proteins to generate hybrid chimeric CARD-CC proteins, even when the sequences are not conserved. A striking unique advantage with the CARD9 backbone seems to be the inhibitory capacity of the C terminal sequence, which is able to auto-inhibit all the other paralogs (Fig. 4A). The chimeric proteins all activate down-stream signaling, with some protein domain transplantations resulting in reduced or inefficient auto-inhibition. This indicates that critical intramolecular interactions that keep the protein in an inactive state represent a significant divergence between the CARD-CC paralogs. Our chimeric CARD9 constructs are rather suitable for high-throughput analyses of several candidate activation mutations upon simple transient transfection in HEK293T cells. As can be seen with many constructs, the CARD9 backbone results in dramatic fold inductions, while the same mutations in the native protein give much weaker effects due to a higher background signal in HEK293T cells. The use of chimeric CARD9 backbones also revealed that some activating mutations act internally on the domain where they are located, which is in contrast to previously reported models where mutations were proposed to disrupt intra-molecular but inter-domain interactions between CARD, LATCH and CC or between CC and the inhibitory region/linker [15,23,56]. Utilization of such functionally conserved but sequence-diverged protein domains is thus a powerful complementary approach to specific site-directed mutagenesis in order to understand the mechanisms and structural constraints for protein function [2,87,88]. For example, there is a high representation of psoriasis-associated CARD14 mutations in the poorly conserved LATCH domain (Fig. 1,4D, 6A). Chimeric CARD9 constructs to study these CARD14 LATCH mutations still need further optimization since only one mutation (L124P) was activating the CARD9-14L chimera. Identification of the required surrounding sequence for a functional chimeric CARD9 CARD14 LATCH psoriasis mutation reporter for mutations at G117 and M119 will also give further structural insight in how CARD14 LATCH gain-of-function mutations require specific interactions with those surrounding sequences. The sequence-swapped chimeric CARD9 construct should also be ideal for mapping of interacting proteins that specifically bind one CARD-CC isoform. An advantage with these constructs is that they will look and act approximately the same except for the sequence that has been swapped, so there should be fewer artifacts from deletion-induced spontaneous activation or other structural changes. Up until now, only naturally occurring activating germ line mutations have been described in CARD11 (causing B cell lymphoma and BENTA) and CARD14 (causing CAPE) within the CARD-CC family [27]. Activating mutations are typically dominant, and will thus usually be less frequent than recessive loss-of-function mutations if detrimental.

By sequence comparisons with CARD11 and screenings of candidate activating natural variants, we found that there are activating CARD9 mutations in the human germline, and public disease association data were used to mine for more candidate activating variants with similar disease associations. The identification of activating germ-line mutations in CARD9 could be done rapidly by a simple overexpression assay thanks to its intrinsically good auto-inhibition (Fig. 3A,B). Using our novel chimeric CARD9 framework, we could also do a similar simple overexpression-based screening to establish that the POAG-associated CARD10 mutant R212H is activating (Fig. 6C-F), demonstrating that there are germ line-inherited activating mutations also in CARD10. To our surprise, the majority of the POAG-associated CARD10 mutations evaluated in our models did not show signs of gain-of-function — or loss-of-function. Since there are multiple independent mutations in CARD10 that are associated to POAG, it is very likely that those mutations are truly causative of the disease. It is however possible that the activation effect of the CARD10^R212H^ mutation is unrelated to the POAG disease phenotype. Due to the low success rate in finding activating CARD10 mutants by screening published disease-associated mutations, future screening for activating natural variants in CARD10 should probably rather be focusing on other predictions, like the AlphaFold computed structural model [15,71,89] or similarity to other activating mutations (Fig. 1). This was also how we identified the activating CARD10^E145K^ variant and how we engineered the stronger artificially activated CARD10^G128D^ and CARD10^L132P^ mutants (Fig. 2E,G,6F). Also, disease associations that are overlapping with phenotypes displayed by carriers of already known activating variants in CARD10 could be very informative, as demonstrated by our identification of the activating CARD10^R153P^ variant based on the disease associations of CARD10^R212H^. The relatively mild phenotype (POAG) officially associated to the activated CARD10^R212H^ is quite surprising considering the broad expression profile of CARD10, and the quite severe diseases that are associated to genetic activation of CARD11 in lymphocytes and CARD14 in keratinocytes. It is possible that the phenotypic effect of CARD10 activating mutations is blunted thanks to the MALT1 protease-mediated negative feedback regulation through CARD10 cleavage [90].

The activating mutations in CARD9 and CARD10 were only picked up in Genebass as a heterozygote and very rare, which indicates that they are (co-)dominant activating mutations. It should however be noted that known dominant CAPE mutations in CARD14 and known dominant BENTA mutations in CARD11 are also absent or as rare as the activating CARD9 or CARD10 mutations in the Genebass database, and thus probably also equally rare in the general population. The dominant nature of the gain-of-function activating mutations allowed us to identify potential novel disease associations with the six activated CARD9 variants and three activated CARD10 variants that we identified in our screening. In contrast, rare recessive loss-of-function mutations are often only causing disease in populations with high levels of endogamy [91], which would have been difficult to pick up in Genebass due to limited ethnic diversity in the UK Biobank dataset [63].

Taken together, we have thus confirmed the existence of germ-line activating mutations in both CARD9 and CARD10, which means that all four CARD-CC paralogs have activating mutations in the human gene pool. Many of the activating variants are in residues conserved between the CARD-CC paralogs. It is quite fascinating how residues that have remained conserved during approximately half a billion years of vertebrate evolution can be mutated and inherited in small fractions of the human population. This high level of conservation does however suggest a very strong selective pressure to maintain control of CARD-CC activation, and we expect aberrant activation of any of the four CARD-CC proteins to be very detrimental and counterselected.

It would be very interesting to investigate the biological effect of expressing the activated CARD9^G111E^ mutation by knock-in mutation of the *Card9* gene in mice. Since it persists in germline in humans, we would expect that such mice would be viable. However, previous studies have shown that a CARD14 mutation (E138A) that is non-lethal and causing psoriasis in humans can be lethal in mice [92–94]. If the endogenous knock-in is more detrimental in mice than in humans, an alternative would be expressing them conditionally in myeloid cells using a *Rosa26*^lox-stop-lox^ system in mice, similar to what has been done previously with conditional transgenic expression of CARD11^L232LI^, CARD11^ΔID^ or CARD14^E138A^ to study their effects in B cells and keratinocytes, respectively [93,95,96]. The knock-in allele could also be expressed conditionally from its native locus, like what has also been done for *Card14*^E138A^ [97].

The functional verification of the activating *Card10*^R212H^ mutation by using knock-in mice would also be highly interesting. Since so many POAG-associated CARD10 mutants did not show any signs of activation, we would expect other phenotypes in addition to POAG in this model. Alternatively, the stronger activated artificial mutant *Card10*^G128D^ or *Card10*^L132P^ knock-in mice would be interesting to generate in order to specifically look at the physiological effects of CARD10 activation without the potential additional off-target association to POAG. The activated *Card10* knock-in mouse model could thus also provide a valuable novel model to investigate the effects of hyperactive CARD10 in other diseases, such as cancer, asthma, liver- and lung fibrosis [98–101]. Some additional Genebass [63] associations for this natural CARD10^R212H^ variant indicate that such phenotypes may be likely. Of special interest in this model will be the analysis of kidney function since this is a common disease association of the activated CARD10^R212H^ and CARD10^R153P^ variants. The availability of public exome data from biobanks with disease associations and other metadata gives unprecedented opportunities for reverse genetics approaches to find novel disease-associated mutations. We can now first establish the function of a natural variant *in silico* and *in vitro* and then determine the phenotypic consequences and relevance of that activity via searches in public disease association databases. It is possible that traditional forward genetics approaches would have been unable to identify the activated CARD9 and CARD10 mutations due to their rarity and associations to complex disease phenotypes.

In conclusion, the here described CARD9 chimeria overexpression approach in HEK293T cells provides an elegant and quick system to perform structure-function analysis of CARD-CC family members and to predict the potential impact of specific mutations or variants on CARD-CC function. However, it is important to also keep in mind some limitations. A major weakness of the system is that several CARD-CC or chimeric proteins are expressed at different levels, complicating their functional comparison. WT CARD9 expression levels are very low, even at plasmid DNA transfection levels, which appears to be at least partially due to the C terminal inhibitory sequence (Fig. 4A). The very low basal activity of CARD9 leads to extremely high fold-inductions of NF-κB activity with different variants, but the interpretation of the results is complicated by highly variable protein expression between active and inactive CARD9 variants (Fig. 3A). In some of our experiments, we even had to overexpose western blots in order to visualize the expression of WT CARD9, which led to either oversaturated bands (Fig. 3B) or visualization of degradation products from other proteins (Fig. 2G, 4F) in the same western blot. This low expression seems to reflect post-transcriptional effects as differences in CARD9 protein expression are still observed after correction for CARD9 mRNA expression levels (Fig. 3C, D). We have therefore chosen to not normalize expression levels by adapting plasmid concentrations since the observed differences in expression may also reflect real functional effects of the different mutations on protein stability or solubility. For example, CARD9 and CARD11 activation is known to be associated with the formation of insoluble CBM filaments [12,33]. Another limitation of our approach is that CARD-CC function is tested in HEK293T cells. In this context, it is important to note that, in contrast to the high NF-κB activity upon CARD14 or CARD14sh expression in HEK293T cells, activity is very low when expressed in the HaCaT keratinocyte cell line, where G117S mutation has a strong activating effect [24]. Although this may also reflect differences in expression levels in HEK293T versus HaCaT cells, it is not unlikely that also the specific cellular context has a strong influence on CARD-CC activity. Nevertheless, even in cases where the CARD-CC isoform and the cellular context match, such as CARD10 which is known to be endogenously expressed in HEK293T cells, transfection of non-chimeric mutant CARD-CC proteins may still not allow to detect activating mutations. This is illustrated by our observation that activating natural variants could not be verified in the intact CARD10 protein, which may be due to too high expression levels, even at low concentrations of transfected plasmid DNA. Taken together, we believe that various CARD-chimeras can be helpful tools for structure/function analyses and to investigate differential activation of CARD-CC variants in initial studies. Follow-up mechanistic studies of a specific and clinically relevant mutation will then be best performed in an as native cellular environment as possible and eventually analyzed for its pathological relevance in a full organism using well-defined knock-in approaches.

## Material and Methods

### Biological material availability

All plasmid constructs used and generated in this study are available via BCCM/GeneCorner (www.genecorner.ugent.be) and can be found back using the catalog (LMBP) numbers.

### Sequence and structure comparisons

Sequence analysis, alignments and primer design was done in UGene [102] installed via BioArchLinux [103]. Protein domain coordinates were collected from Uniprot [62], and the coordinates were subsequently verified visually using the Alphafold [89] database at EBI (https://alphafold.ebi.ac.uk/). In order to evaluate potential multimeric forms of the CARD-CC protein structures, we ran Alphafold-Multimer [104] on Vlaams Supercomputer Centrum (VCS) (https://www.vscentrum.be/) for CARD9, CARD10, CARD11 and CARD14 using the default settings. All CARD-CC proteins ended up with a better dimeric structure than the previous monomeric prediction available at EBI. The resulting protein structure model (pdb) files (selected based on lowest prediction number and relaxed) were visualized and analyzed in Mol* (https://molstar.org/viewer/) [105]. Mol* was also used to export the raster graphics snapshots used in Figure 2A. For a higher-quality view of the generated dimeric structural models, the pdb files, confidence values and details about the modelling procedure were uploaded to the ModelArchive (https://modelarchive.org) database with the accession codes ma-5s006 (CARD9), ma-hmm6l (CARD10), ma-tzgev (CARD11) and ma-yk1bv (CARD14). We also modeled the hypothetical CARD10/CARD14 heterodimer [10] with the accession code ma-11i6w. The overlap between the calculated structural model of CARD9 and the published experimentally determined dimeric N terminal structure of CARD9 (pdb 6N2M) [15] was investigated by superimposing the structures in Mol*. The start of the CARD10 PDZ domain was identified by graphical investigation in Alphafold structural models only since it was not annotated in Uniprot.

### Cloning hybrid chimeric CARD-CC fusion proteins and other mutants

In order to obtain completely scar-free fusion proteins, we employed NEBuilder (New England Biolabs) to fuse PCR products. PCR fragments were amplified using Phusion DNA polymerase (New England Biolabs) using the generic protocol and 30 s/Kb extension. Our general strategy was to amplify half of the plasmid starting from the middle of the ampicilin resistance gene and the part of the CARD-CC protein that we wanted to fuse, and then fuse the two halves of the plasmid using NEBuilder. For an overview of primers used, see (table 2), and for an overview of constructs generated, see (table 3). Original templates used for the PCR were pCDNA3-F-hCARD9 (LMBP 09609), pCDNA3-F-mCard10 (LMBP 10187), pCD-F-bio-mCard11 (LMBP 10717), pCD-F-bio-mCard11-L232LI (LMBP 07889), pCDNA3-F-hCARD14 (LMBP 09869), pCDNA3-F-hCARD14-E138A (LMBP 09860). The most optimal CARD14-derived chimeric CARD9 variants (C9-14L and C9-14CC) were utilized to quantitatively investigate several known psoriasis-associated mutants (table 4), which were introduced by amplification of the plasmid with the indicated mutation primers followed by DpnI (Promega) digestion. The CARD10 R404W, R420C, ΔK272-E273 and ΔK272-V277 (table 4) mutations [39–41] were also tested in the C9-10CC backbone using the same strategy. The substitutions in mutated proteins were named after the human coordinates to easily cross-reference with human genetic data, even if the mutation was generated on a mouse sequence. For future simplification of high-throughput mutagenesis, C9-14L (LMBP 13438), C9-14CC (LMBP 13437) and C9-10CC (LMBP 13439) were also re-cloned into the pICOz [106] (LMBP 11103) backbone. All the generated clones were verified using Sanger sequencing (Mix2Seq ON, Eurofins Genomics).

**Table 2.**
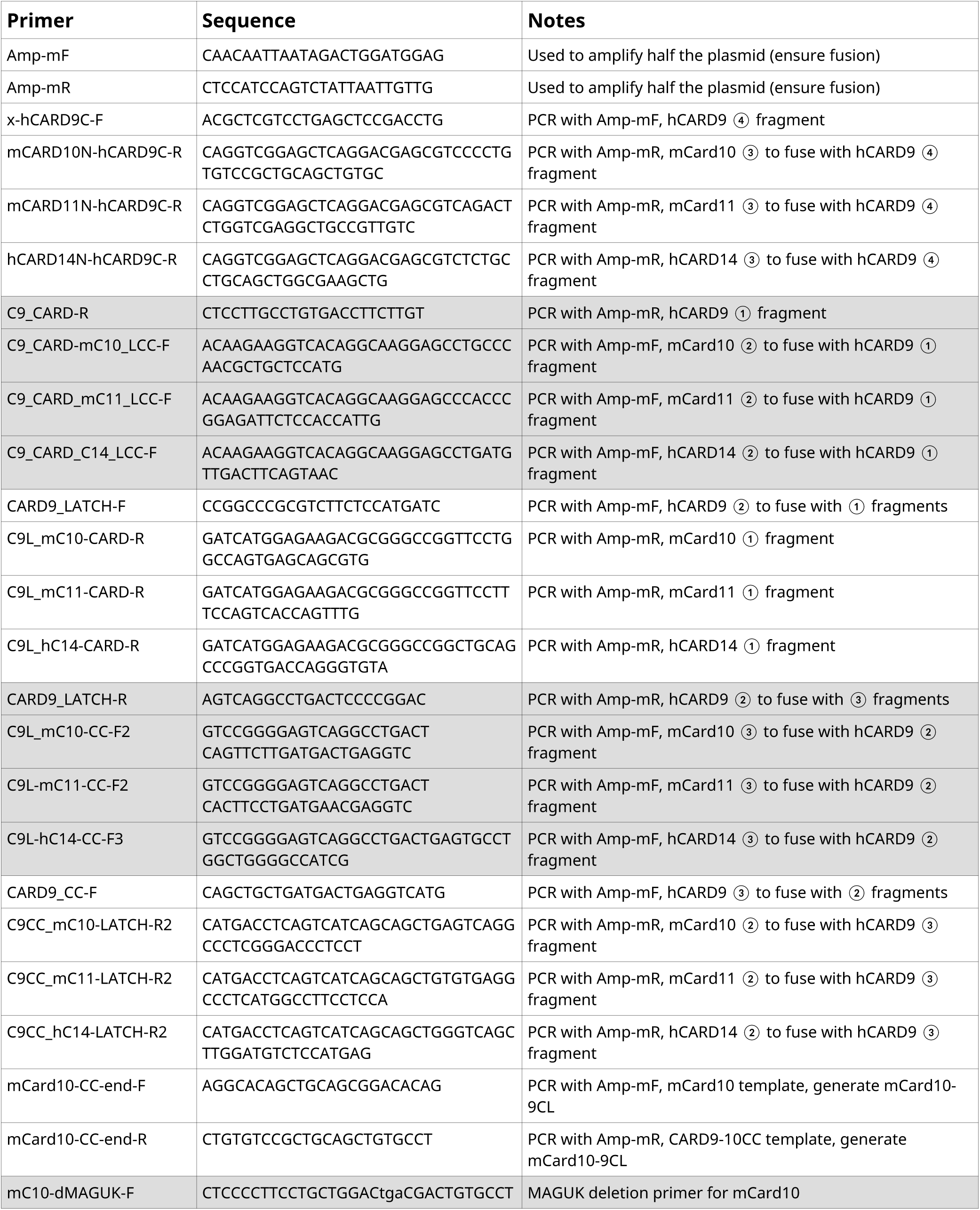
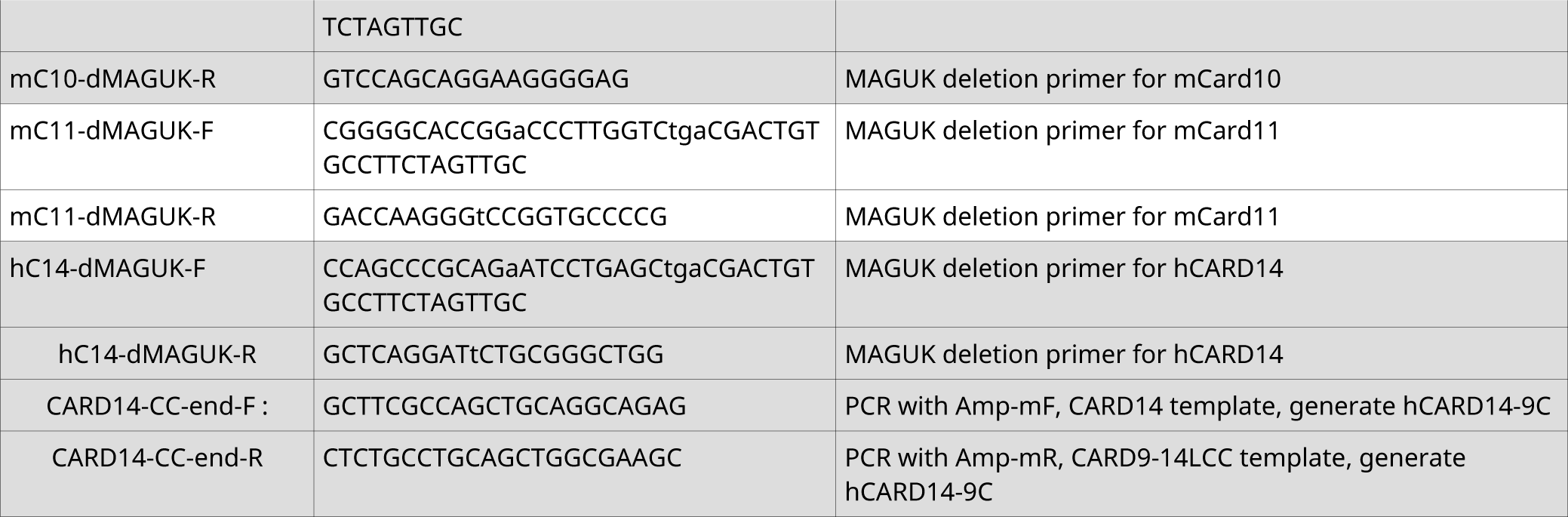
Primers used for fusion CARD-CC proteins and mutants.

**Table 3.**
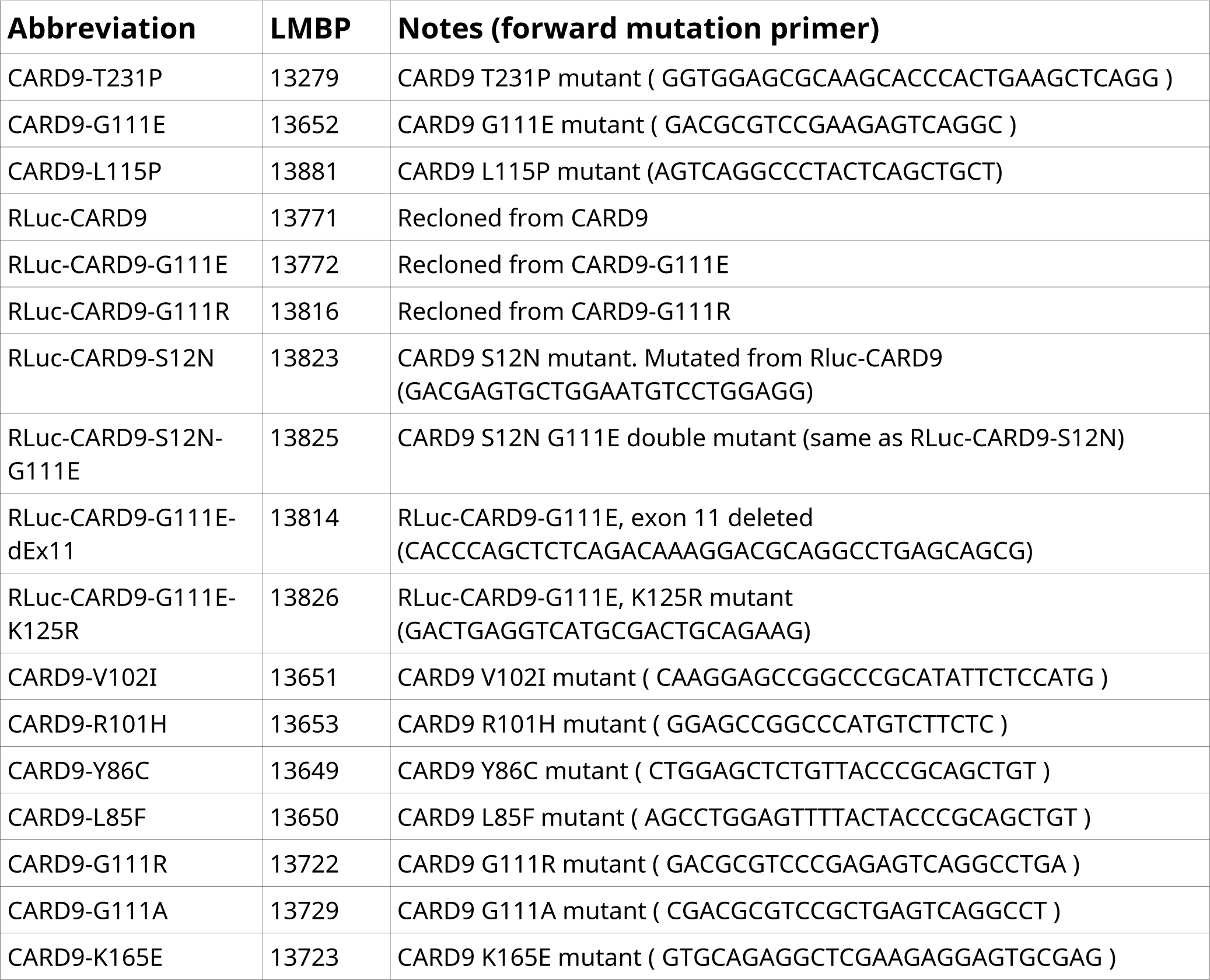
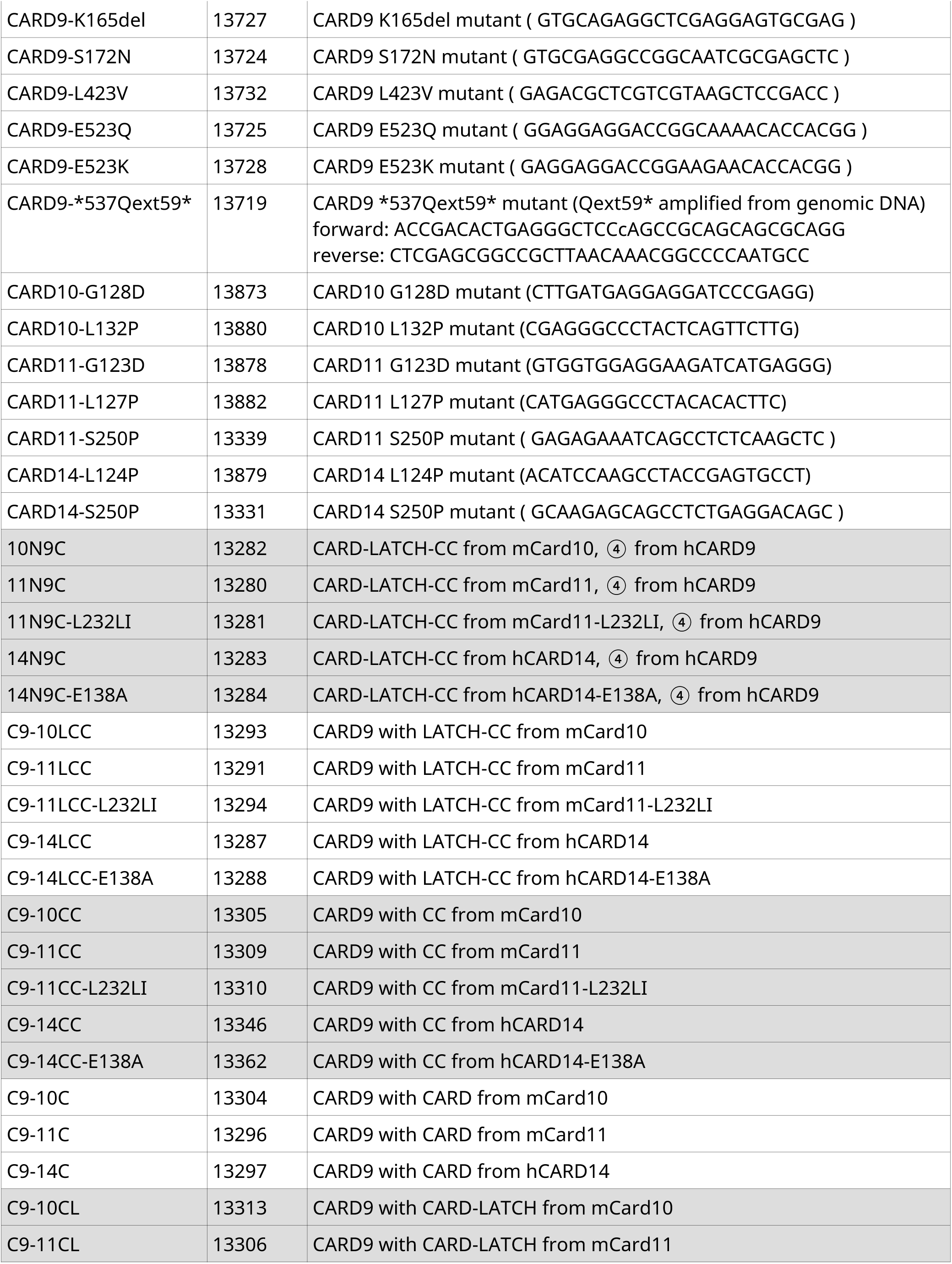
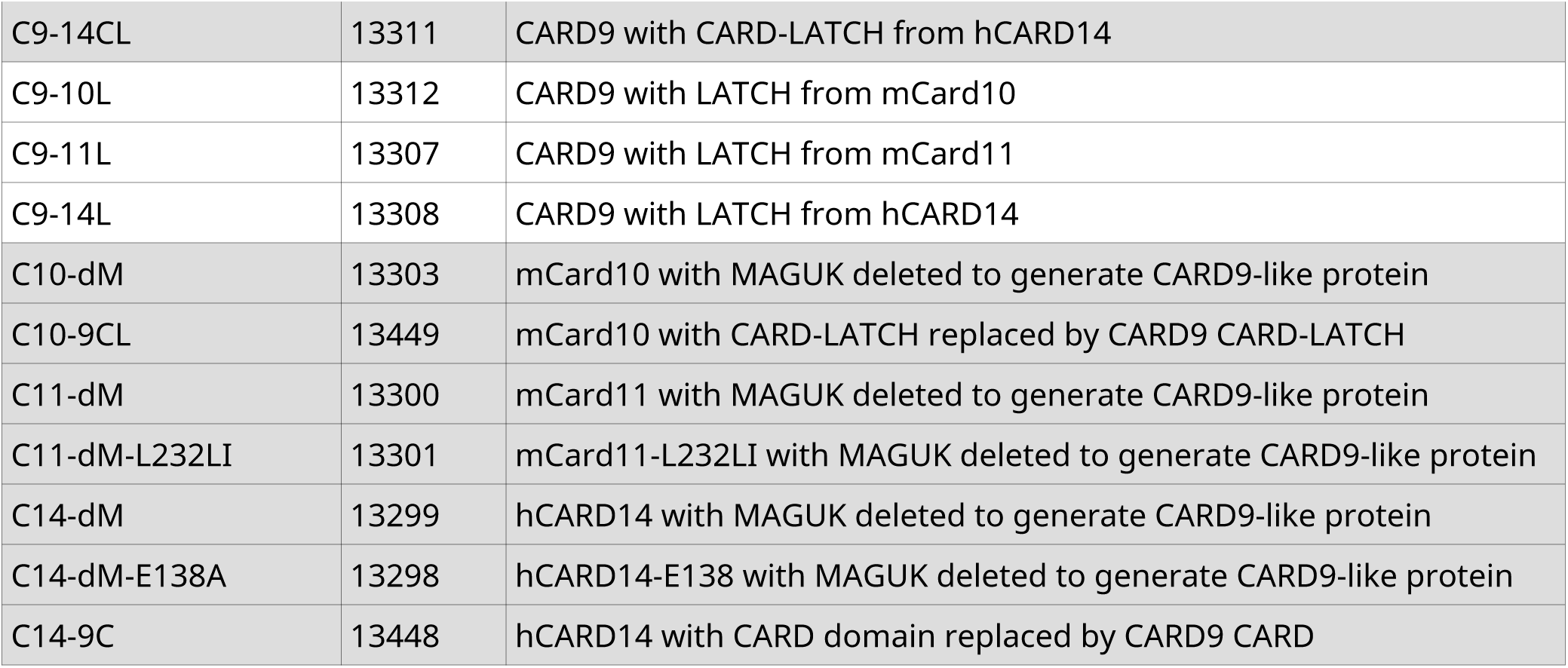
Overview of constructs generated in this study. Further derivative constructs made in this study can be found in table 4.

**Table 4.**
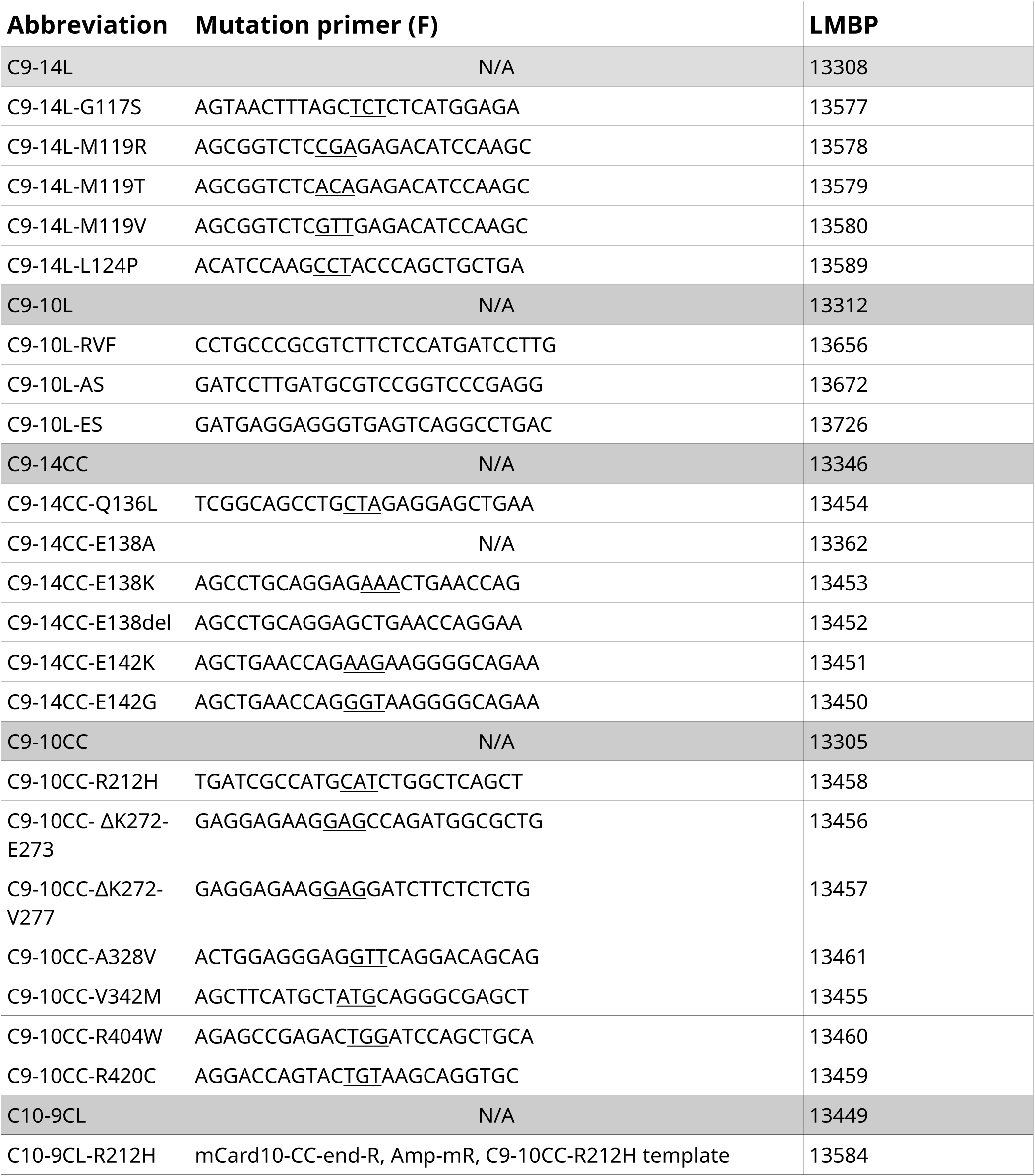
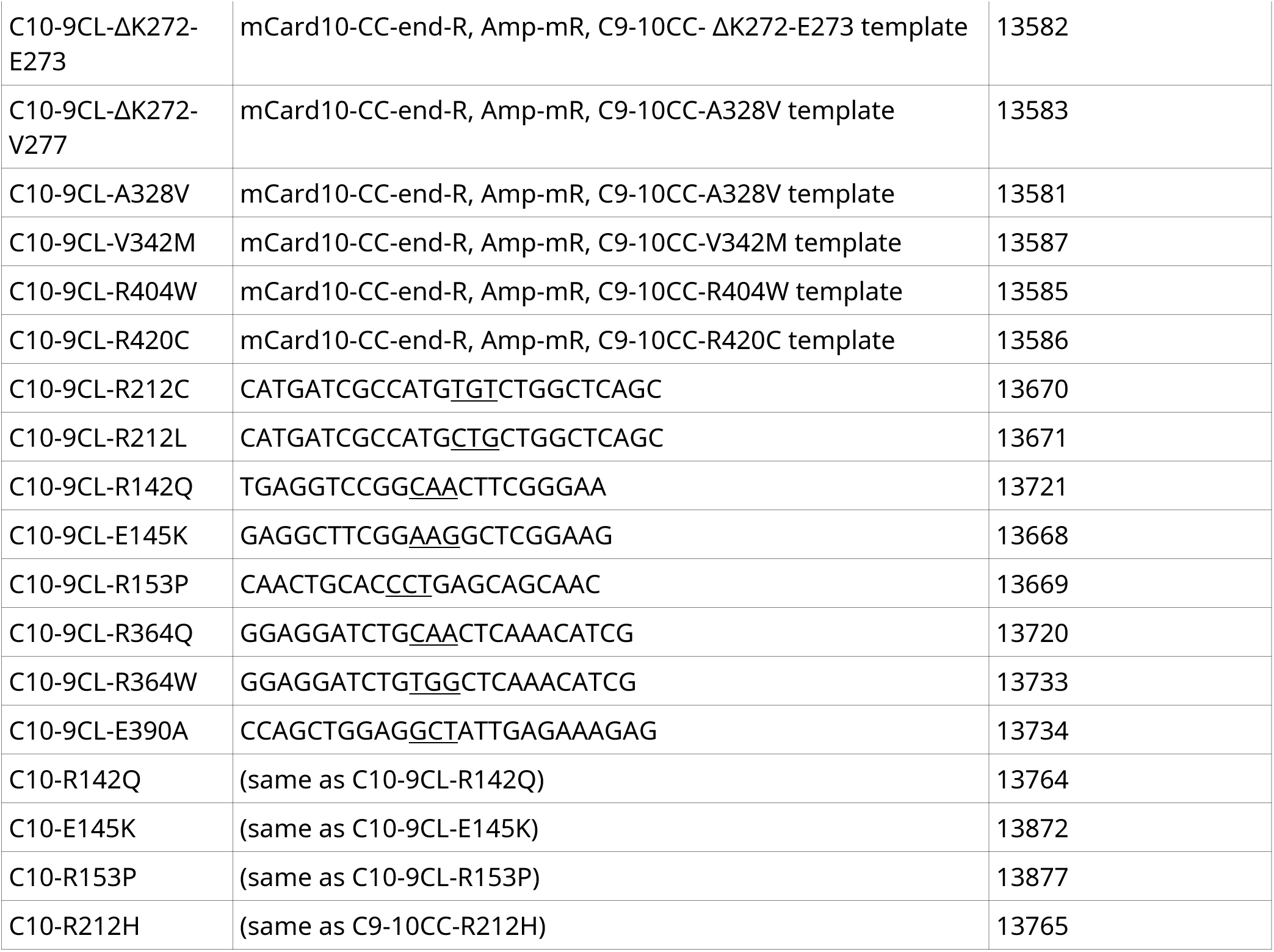
Clones of published LATCH-CC-domain mutations in CARD10 and CARD14 using the minimal chimeric CARD9 backbones C9-14L, C9-14CC, C9-10L, C9-10CC and the good auto-inhibited mouse CARD10 backbone C10-9CL.

### Other CARD-CC protein plasmid resources

Already cloned [2,33] constructs used in this study are pCDNA3-Flag-hCARD9-L213LI (LMBP 10178), pCDNA3-Flag-hCARD9ΔC (“9N”, LMBP 10457), pCDNA3-F-mCard10ΔC (“10N”, LMBP 10459), pCD-F-bio-mCard11ΔC (“11N”, LMBP 10458) and pCDNA3-F-hCARD14ΔC (“14N”, LMBP 10460).

### NF-κB luciferase analysis

For each set-up, we made a master mix of 200 ng/μg CARD-CC expression clone, 100 ng/μg NF-κB-dependent luciferase expression vector (LMBP 3249) [107], 100 ng/μg actin-promoter driven β-galactosidase (LacZ) expression plasmid (LMBP 4341) [108] and 600 ng/μg empty vector (LMBP 2453). 1 μg master mix was used to transfect 4 wells in a 24-well plate of HEK293T cells (200 000 cells/well) using the calcium phosphate transfection method. Statistical tests for differences in the luciferase results were done using Student’s t-test [109] and the probability levels of non-rejection of the null hypothesis that two values are equal are abbreviated as * = p ≤ 0.05, ** = p ≤ 0.01, *** = p ≤ 0.001 and **** = p ≤ 0.0001.

### Western blot analysis

To ensure that the chimeric CARD-CC proteins are expressed, we lysed a set of transfected cells in Laemmli buffer [110], ran the samples on SDS-PAGE, and subsequently blotted the proteins on nitrocellulose membranes by wet blotting. Since all the cloned CARD-CC proteins are N-terminally flag-tagged, we developed the blots using anti-Flag (clone M2, F-3165, Sigma) and detected them on an Amersham Imager 600. As loading control, we developed for anti-actin (sc-47778, Santa cruz).

## Abbreviations

BCL10: B-cell lymphoma 10
BENTA: B cell expansion with NF-κB and T cell anergy
CARD: caspase recruitment domain
CAPE: *CARD14* – Associated Papulosquamous Eruption
CARMA: CARD-containing MAGUK protein
CBM: CARD-CC/BCL10/MALT1 complex
CC: coiled-coil domain
HEK293T cells: Human embryonic kidney 293T cells
MAGUK: membrane-associated guanylate kinases
IBS: Irritable bowel syndrome
MALT1: mucosa associated lymphoid tissue lymphoma translocation 1
MUSCLE: MUltiple Sequence Comparison by Log- Expectation
POAG: Primary open angle glaucoma
RMSD: root mean square deviation
WT: wild-type

## Author contributions

Y.D, Ji.S, F.V.G and Je.S performed experiments. Je.S. conceptualized, planned and wrote the manuscript. Je.S and R.B. supervised the project.

## Acknowledgements

Research in the authors’ lab is supported by a VIB Grand Challenges grant (GC01-C01), the Fund for Scientific Research Flanders (FWO;3G086521), an Excellence of Science (EOS) research grant (3G0I1422), Ghent University Concerted Research Actions (GOA; 01G00419). F.V.G. holds a PhD fellowship strategic basic research from the FWO. Je.S. holds a doctoral assistant mandate and assistant professorship from Ghent University. Disease associations have been conducted using summary statistics generated from the UK Biobank resource (under application 26041 and 48511).

---

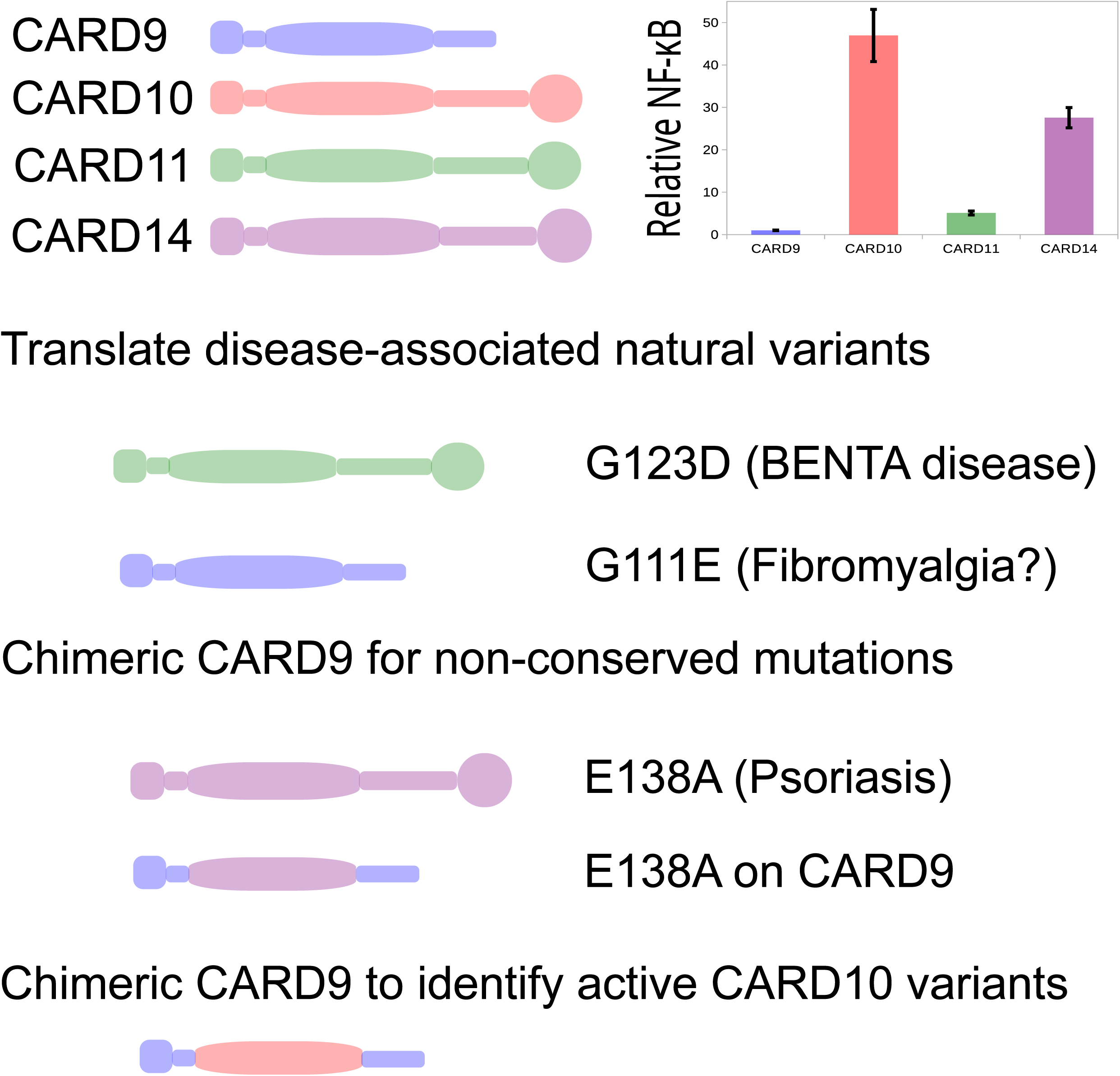

